# Extracellular vesicles and their RNA cargo facilitate bidirectional cross-kingdom communication between human and bacterial cells

**DOI:** 10.1101/2025.07.10.664124

**Authors:** Laura Gröger, Shusruto Rishik, Nicole Ludwig, Amila Beganovic, Marcus Koch, Stefanie Rheinheimer, Martin Hart, Petra König, Tabea Trampert, Pascal Paul, Annette Boese, Claus-Michael Lehr, Sören L. Becker, Gregor Fuhrmann, Andreas Keller, Eckart Meese

**Affiliations:** Saarland University (USAAR), Department of Human Genetics, 66421 Homburg, Germany; Saarland University (USAAR), Chair for Clinical Bioinformatics, Center for Bioinformatics, 66123 Saarbrücken, Germany; INM - Leibniz Institute for New Materials, 66123 Saarbrücken, Germany; Saarland University of Applied Sciences, Institute for Physical Process Technology, 66117 Saarbrücken, Germany; Saarland University (USAAR), Center for Human and Molecular Biology (ZHMB); Helmholtz Institute for Pharmaceutical Research Saarland (HIPS), 66123 Saarbrücken, Germany; Saarland University (USAAR), Department of Pharmacy, 66123 Saarbrücken, Germany; Saarland University (USAAR), Institute of Medical Microbiology and Hygiene, 66421 Homburg, Germany; Department of Biology, Pharmaceutical Biology, Friedrich-Alexander-University (FAU) Erlangen-Nürnberg, 91058 Erlangen, Germany; Saarland University (USAAR), PharmaScienceHub (PSH), 66123 Saarbrücken, Germany

**Keywords:** Extracellular vesicles, EV, Communication, Cross-Kingdom, Bacteria, miRNA

## Abstract

While extracellular vesicles (EVs) are established mediators of intra-species signaling, their role as active participants in cross-kingdom communication remains incompletely understood. Here, we reveal that human colon cells and both Gram-positive and Gram-negative gut bacteria engage in species-specific, EV-mediated molecular dialogue, driven in part by RNA cargo. We show that bacterial EVs (BEVs) induce distinct transcriptomic responses in human cells, and that BEV-RNA independently causes similar effects. Conversely, we demonstrate that human EVs and highly abundant miR-192-5p are differentially internalized by bacteria, affecting their physiology. Our findings support a conceptual model in which EVs function as directional messengers that shape host-microbiome interactions. This study introduces a framework for understanding EVs as cross-kingdom regulators and underscores the importance of tailored, context-specific analyses for understanding the scope of EV-mediated interactions in microbiome-host homeostasis and disease.

**Highlights:**

[1] *L. casei*, *E. faecalis* and *P. mirabilis* produce BEVs that are internalized by Caco-2 cells at different rates. BEVs produced by *L. casei* have a positive influence on the viability of Caco-2 cells. Incubation of Caco-2 cells with BEVs leads to changes in the gene expression of immune-response-related genes.
[2] BEVs carry RNAs and the type of RNA cargo varies significantly between the BEVs from the different bacteria. Comparison of Caco-2 gene deregulation between BEVs and transfection of RNA isolated from BEV highlights component-specific effects.
[3] Caco-2 EVs are taken up by *E. faecalis* and influence their growth. MiRNA-192-5p can be frequently detected in EVs from Caco-2 cells. Synthetic miR-192-5p is internalized by *P. mirabilis* and the ability to take up human miRNAs by *L. casei* and *E. faecalis* can be increased by packaging of the miRNA in artificial liposomes.

## Introduction

Communication between cells is essential for building and maintaining complex organization in higher organisms. While this communication can occur through direct cell-to-cell contact or by secretion of signaling molecules into extracellular fluids to reach distant recipient cells [1], researchers have proposed that extracellular vesicles (EVs) act as versatile and conserved mediators of cell-to-cell communication [2]. EVs are small membrane-bound particles naturally released into the extracellular space. They are produced by most eukaryotes and prokaryotes and transport various molecules such as proteins, lipids, and nucleic acids [3]. Among other classes of RNAs, EVs also carry microRNAs (miRNAs), which are known to act as important posttranscriptional regulators of gene expression [4, 5].

While EVs are well established as participants in intra-species communication, more recent evidence suggests that they also actively facilitate cross-kingdom interactions [2]. Various studies indicate that bacterial EVs (BEVs), including membrane vesicles (MVs) produced by Gram-positive bacteria, as well as outer membrane vesicles (OMVs) produced by Gram-negative bacteria, can be transferred to eukaryotic cells and modulate host gene expression [6–9], with bacterial RNA cargo playing a significant role in this modulation [10–12]. Despite broad agreement on the relevance of BEVs in bacterial-mammalian cross-kingdom communication, key mechanistic details remain debated, and the impact of bacterial RNAs on gene regulation in mammalian cells remains largely unknown. This gap in knowledge is especially relevant given the association of BEVs with human diseases such as inflammatory bowel disease [13]. It is therefore crucial to gain deeper insights into these communication process at the highest possible resolution. Furthermore, even in diseases not directly associated with the gut – such as Parkinson’s disease (PD) – changes in the gut microbiome composition are frequently observed [14, 15], raising the possibility that BEVs may contribute to disease development and progression. Conversely, several studies showed that human and murine miRNAs can be taken up by bacterial cells and can affect their growth and gene expression [16–20]. However, the impact of host-derived EVs and their RNA cargo on gut bacteria remains similarly underexplored.

Here, we address this gap by proposing that EVs and their RNA cargo mediate a bidirectional, species-specific molecular dialogue between human colon cells and different gut bacteria. Using a model system comprising Gram-positive and Gram-negative bacteria of the order Lactobacillales and Enterobacterales [21], which have been frequently associated with dysbiosis in PD, we systematically dissect how BEVs and BEV-RNA affect the host transcriptome. Conversely, we explore how host EVs influence bacterial growth and whether bacteria are able to internalize human miR-192-5p. Rather than treating EV exchange as a passive or incidental process, our findings support the conceptual framework in which EVs act as targeted messengers that facilitate structured cross-kingdom communication. This work not only defines the molecular specificity of such interactions but also reframes EVs as active participants in host-microbiome regulation with potential implications for health and disease.

## Results

### Lacticaseibacillus casei, Enterococcus faecalis and Proteus mirabilis produce BEVs that are taken up by Caco-2 cells

To study the effect of BEVs on eukaryotic recipient cells, we first isolated BEVs from bacterial cultures (**Figure S1**) using ultracentrifugation and size-exclusion chromatography (SEC), and characterized them by nanoparticle tracking analysis (NTA) and cryo-transmission electron microscopy (Cryo-TEM). Looking at the BEVs from each bacterium individually, we found that *Lacticaseibacillus casei* (*L. casei*) MVs had a mean size of 140.0 (± 15.5) nm (**Figure 1A**), *Enterococcus faecalis* (*E. faecalis*) MVs of 117.1 (± 7.5) nm (**Figure 1B**), and *Proteus mirabilis* (*P. mirabilis*) OMVs of 115.2 (± 7.3) nm (**Figure 1C**). Additionally, we found that *L. casei* MVs had a particle concentration of 2.1x10^12^ (± 6.4x10^11^) particles/ml, while *E. faecalis* MVs had a concentration of 8.5x10^10^ (± 8.2x10^9^) particles/ml, and *P. mirabilis* OMVs of 6.4x10^10^ (± 1.0x10^10^) particles/ml. Cryo-TEM images of BEVs showed round-shaped particles (**Figure 1D-F**). Since OMVs from Gram-negative bacteria harbor LPS [22], we furthermore quantified the LPS concentration of *P. mirabilis* OMVs, demonstrating that OMVs contained 1.55 (± 0.15) ng/ml LPS (**Figure S2A**).

**Figure 1:**
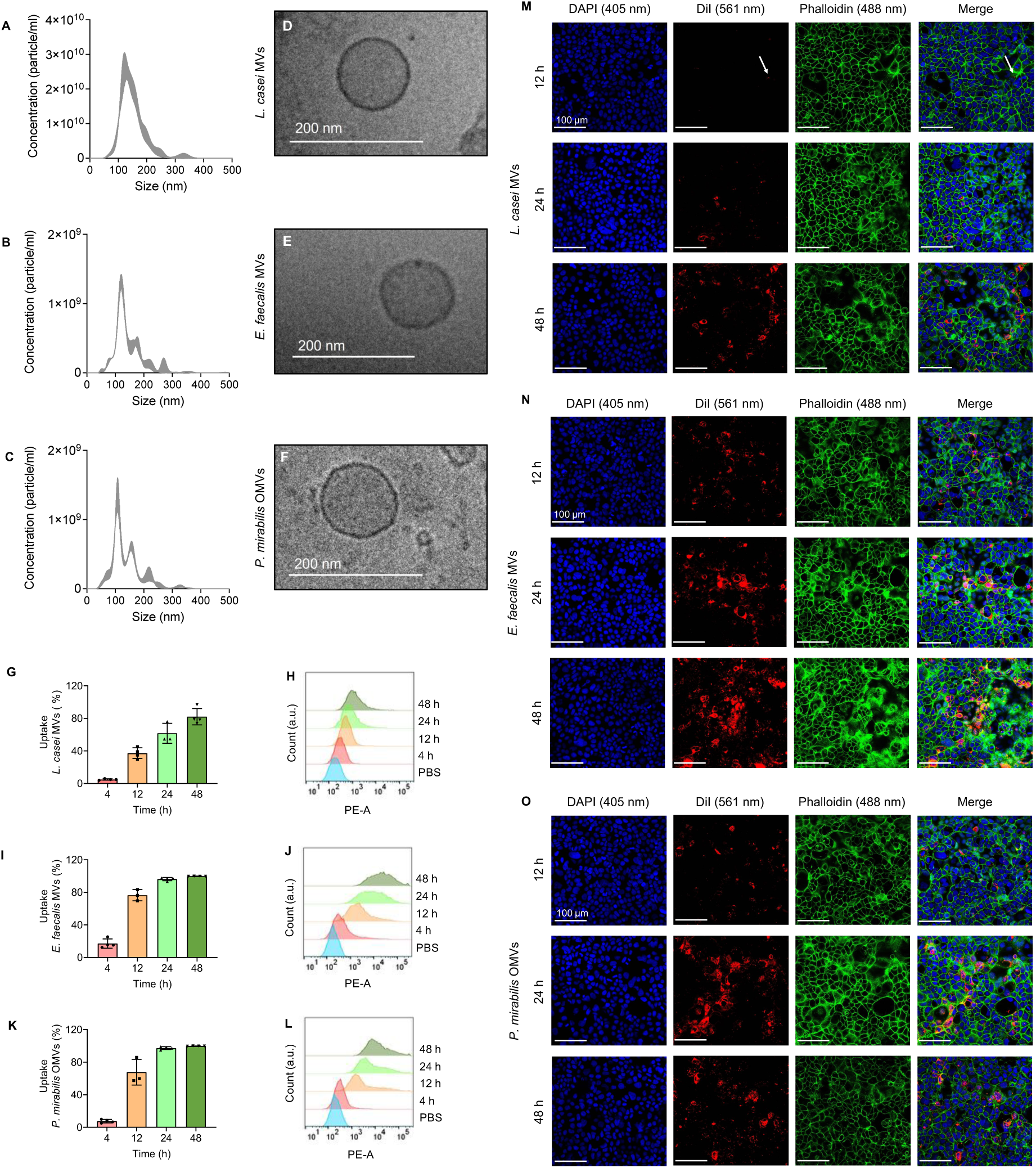
Uptake of BEVs from *L. casei*, *E. faecalis* and *P. mirabilis* by Caco-2 cells. (**A-E**) Characterization of BEVs. Typical size distribution of purified BEVs of *L. casei* MVs (**A**), *E. faecalis* MVs (**B**), and *P. mirabilis* OMVs (**C**) measured by nanoparticle tracking analysis. Representative cryo-TEM images of *L. casei* MVs (**D**), *E. faecalis* MVs (**E**), and *P. mirabilis* OMVs (**F**) showed round shaped particles. (**G-L**) Flow cytometry analysis of the uptake of fluorescence-labeled BEVs in Caco-2 cells. Percentage of PE (Phycoerythrin)-positive cells and representative histograms compared to the control (PBS) after 4 h, 12 h, 24 h, or 48 h incubation with the DiI-labeled BEVs of *L. casei* (**G and H**), *E. faecalis* (**I and J**), or *P. mirabilis* (**K and L**). Results are presented as mean ± standard deviation (SD) from 3-4 independent experiments. (**M-O**) Visualization of the internalization of BEVs into Caco-2 cells using CLSM. Caco-2 cells were incubated for 12 h, 24 h, or 48 h with DiI-labeled *L. casei* MVs (**M**), *E. faecalis* MVs (**N**), or *P. mirabilis* OMVs (**O**). After the incubation, the cells were washed, fixed, and permeabilized. F-Actin was stained with phalloidin, and the cell nuclei were stained with DAPI. DAPI was visualized using a diode laser at 405 nm (blue), phalloidin with an argon laser at 488 nm (green), and DiI with a DPSS laser at 561 nm (red). The white arrow indicates an area where weak signals for DiI-labeled *L. casei* MVs were detected. Scale bar: 100 µm.

Next, we asked whether Caco-2 cells can take up BEVs from *L. casei*, *E. faecalis,* and *P. mirabilis.* Using flow cytometry, we found that out of all three BEV types, Caco-2 cells take up *L. casei* MVs with the slowest rate, displaying a maximum of 82.1 (± 10.1) % fluorescent cells after 48 h (**Figure 1G and 1H**). Remarkably, the human cells take up BEVs from *E. faecalis* (**Figure 1I and 1J**) and *P. mirabilis* (**Figure 1K and 1L**) at significantly faster rates. To validate the actual internalization of DiI-labeled BEVs we used confocal laser scanning microscopy (CLSM). In accordance with the previous results, we observed little accumulation of *L. casei* MVs in the cells after 12 h, which increased with longer incubation time. For cells incubated with *E. faecalis* MVs or *P. mirabilis* OMVs, we detected a strong accumulation of BEVs within the cells already after 12 h of incubation, which again increased with extended incubation time (**Figure 1M-O**).

### BEVs contain different RNA amounts and species and incubation or transfection of Caco-2 cells with BEVs, or BEV-RNA display no cytotoxic effects

To investigate whether the exposure of Caco-2 cells with BEVs affects the cell viability, we incubated the cells with different concentrations of BEVs for 24 h or 48 h and performed viability and cytotoxicity assays. Indeed, none of the BEVs showed a negative impact on the viability or increased cytotoxicity towards the Caco-2 cells at any of the concentrations tested (**Figure 2A-F**). Interestingly, the incubation of Caco-2 cells with *L. casei* MVs even increased the viability significantly in comparison to the control (**Figure 2A**). This effect was not visible for the other two tested BEVs (**Figure 2B and 2C**). In addition, the viability of Caco-2 cells remained unaffected when incubated with LPS from *P. mirabilis* alone (**Figure S2B and S2C**).

**Figure 2:**
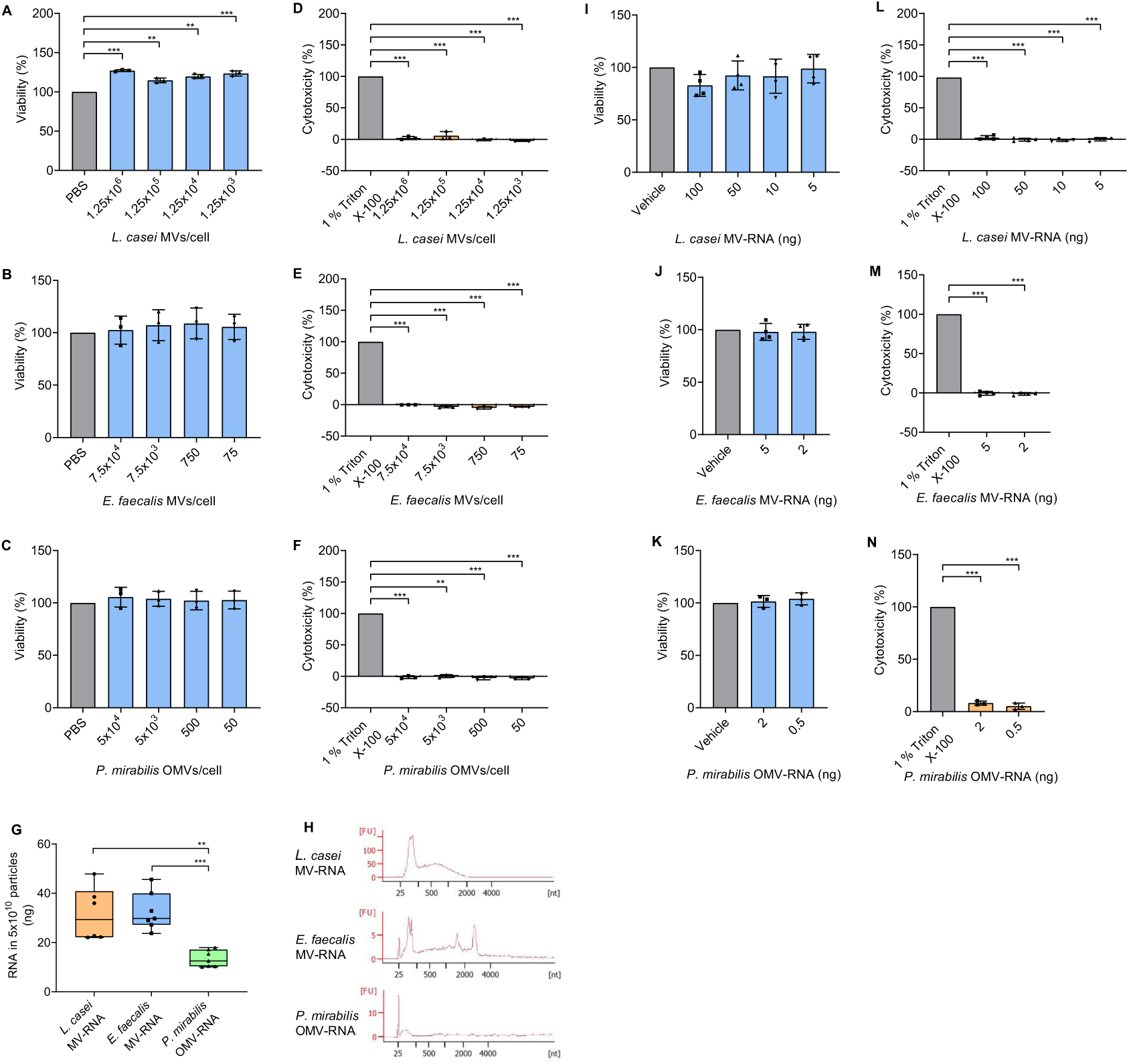
Effects of BEVs and BEV-RNA on Caco-2 cell viability. (**A-F**) Measurement of cell viability and cytotoxicity after incubation of Caco-2 cells with different concentrations of *L. casei* MVs (**A and D**), *E. faecalis* MVs (**B and E**), or *P. mirabilis* OMVs (**C and F**) for 24 h (*E. faecalis* MVs or *P. mirabilis* OMVs) or 48 h (*L. casei* MVs). 1 % Triton X-100 was used as dead control, whereas PBS was used as live control. Results are presented as mean ± SD from 3-4 independent experiments. (**G and H**) Analysis of BEV-RNA. (**G**) Displayed is the amount of RNA present in 5x10^10^ vesicles in box-whisker plots. Results shown are from 6-7 independent experiments. (**H**) Representative electropherograms show size distribution of *L. casei* MV-RNA, *E. faecalis* MV-RNA, and *P. mirabilis* OMV-RNA analyzed using Agilent RNA 6000 Pico Chip. (**I-N**) Measurement of cell viability and cytotoxicity after transfection of Caco-2 cells with different amounts of *L. casei* MV-RNA (**I and L**), *E. faecalis* MV-RNA (**J and M**), or *P. mirabilis* OMV-RNA (**K and N**) for 24 h (*E. faecalis* MV-RNA or *P. mirabilis* OMV-RNA) or 48 h (*L. casei* MV-RNA). 1 % Triton X-100 was used as dead control, whereas Lipofectamine™ 3000 transfection reagent mixed with nuclease-free water (vehicle) was used as live control. Results are presented as mean ± SD from 3-4 independent experiments. For all experiments, a two-tailed unpaired t-test was conducted to determine statistical significance (p < 0.01 **, p < 0.001 ***).

Since BEV-RNA can modulate the gene expression of host cells [10–12], we asked whether this applies also for RNA from *L. casei* MVs, *E. faecalis* MVs or *P. mirabilis* OMVs. We thus isolated and quantified the BEV-RNA content. The overall amount of RNA present in a normalized number of 5x10^10^ particles was comparable between *L. casei* MVs with 31.55 (± 10.86) ng and *E. faecalis* MVs with 32.57 (± 7.65) ng. In contrast, *P. mirabilis* OMVs carried significantly less RNAs with 13.41 (± 3.38) ng RNA in the same number of particles (**Figure 2G**). We further characterized the different RNA species isolated from the BEVs, speculating on differential cargo with respect to long- and short RNAs. Interestingly, we found a high proportion of small RNAs (< 200 nts) but also some larger RNAs of up to 2,000 nts in MVs from *L. casei* (**Figure 2H**). *E. faecalis* MVs also contained many small RNAs, some larger RNA fragments, and ribosomal RNAs. Of note, the latter RNA type was exclusively present in the *E. faecalis* MVs. Most remarkably, *P. mirabilis* OMVs distinguished clearly with respect to their RNA cargo since these OMVs transport exclusively small RNAs. In summary, our results demonstrate that BEVs isolated from different bacteria exhibit diverse RNA cargo. Recently, it has been reported that LPS can co-purify during RNA isolation from OMV [23]. Thus, we also quantified the LPS concentration in the RNA and found that *P. mirabilis* OMV-RNA contained 9.11 (± 0.25) ng/ml LPS. (**Figure S2D)**.

To test whether the transfection of BEV-RNA using artificial liposomes affects the viability of Caco-2 cells, we transfected the cells with different amounts of BEV-RNA for 24 h or 48 h and performed viability and cytotoxicity assays. The amounts tested were chosen based on the amount of RNA that we were able to isolate from the BEVs and that could potentially be delivered into the recipient cells. We again observed stable cell viability and no increased cytotoxicity towards the Caco-2 cells at any of the tested amounts of BEV-RNA (**Figure 2I-N**). We did also not observe any impact on the viability when transfecting the Caco-2 cells with *P. mirabilis* LPS alone (**Figure S2E and S2F**).

### The transcriptome of Caco-2 cells changes in response to incubation with BEVs or transfection with BEV-RNA

To examine changes in the transcriptome of Caco-2 cells due to exposure with BEVs or RNA from BEVs, we performed RNA-seq. We incubated Caco-2 cells with the different BEVs or transfected them with BEV-RNA for 10 h, 24 h, and 48 h exclusively for *L. casei* MVs and MV-RNA due to the slower uptake rate. Based on our previous results, we optimized the particles/cell ratio to ensure sufficient BEV internalization without compromising cell viability. BEV-RNA used for transfection of Caco-2 cells was always isolated from BEVs that were used for the same experiment (**Figure 3A**).

**Figure 3:**
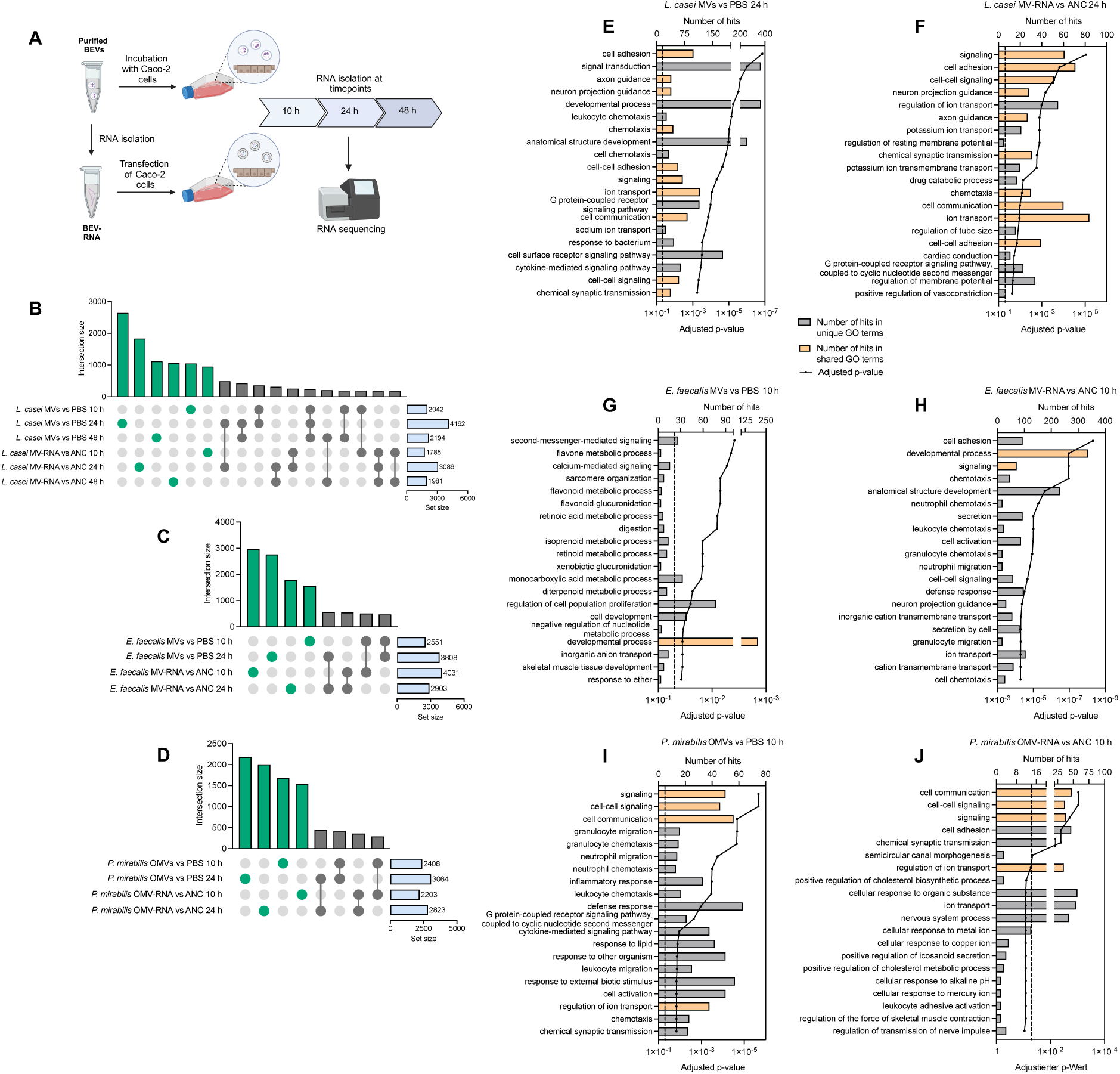
Changes in the gene expression of Caco-2 cells after incubation with BEVs or transfection with BEV-RNA. **(A)** Illustration of experimental workflow during the analysis of the transcriptome (Created with BioRender). Caco-2 cells have been incubated with BEVs (*L. casei* MVs: 1.5x10^11^ particle, *E. faecalis* MVs: 7.5x10^9^ particle, *P. mirabilis* OMVs: 6.0x10^9^ particle) or transfected with BEV-RNA (*L. casei* MV-RNA: 100 ng, *E. faecalis* MV-RNA: 5 ng, *P. mirabilis* OMV-RNA: 2 ng) for 10 h, 24 h, or 48 h. (**B-D**) Upset plots highlight the number of differentially expressed genes between the different time points and treatments. (**E-J**) Differentially expressed genes were used to perform an over-representation analysis (ORA) using GeneTrail 3.2. Shown are the top 20 Gene Ontology (GO) - biological process in which the differentially expressed genes were enriched in. The dashed line marks the significance level of p = 0.05. Biological processes that showed an enrichment after incubation with BEVs and transfection of BEV-RNA derived from the same bacteria at the same timepoint are highlighted in orange.

Across 105 total samples we had 3-4 samples for each timepoint under each condition. We sequenced a median of 124 million reads (IQR of 54 million reads) of which a median of 98.2 % were mapped. After filtering we obtained a median of 107 million reads (IQR of 48 million reads). From these we assigned a median of 4.9 million reads (IQR of 2.1 million reads). After quantification we detected a median of 36,175 genes (IQR of 1086) across the conditions with a total of 46,395 genes detected across all conditions (**Table S1**).

Following incubation of Caco-2 cells with BEVs, we found that most of the genes, including protein-coding genes, pseudogenes, and non-coding RNAs like miRNAs, were altered after 24 h incubation (**Figure S3A-C**). Interestingly, after 48 h incubation of Caco-2 cells with MVs from *L. casei* the total number of differentially expressed genes decreased, even though uptake analysis showed a continuous internalization of the MVs. When transfecting the Caco-2 cells with *L. casei* MV-RNA and *P. mirabilis* OMV-RNA, we again observed that most genes were altered after 24 h. In contrast, we found more genes to be altered in cells transfected with *E. faecalis* MV-RNA after 10 h compared to 24 h (**Figure S3D-F**). While we see a pattern in the overall deregulation after the different treatments, this itself does not tell us if the same genes are taking part in the observed deregulation. Therefore, we investigated which genes that were altered after BEV and BEV-RNA treatments were shared within and across timepoints. We detected a large number of genes that were uniquely deregulated at each timepoint and condition. However, we further observed a smaller gene set showing deregulation at several timepoints and different treatments. This leads to the suggestion that even though the BEVs and BEV-RNA seem to cause a rather short acute deregulation in the expression of specific genes, there are similar events happening which cause a change in the expression of similar genes (**Figure 3B-D**).

To identify specific pathways with which the deregulated genes are associated with, we performed an over-representation analysis (ORA) using GeneTrail 3.2 [24]. For cells incubated with *L. casei* MVs for 10 h and 24 h, we found that the deregulated genes play a pivotal role in a variety of different biological processes like inflammatory response, chemotaxis, and cell adhesion (**Figure 3E, Figure S4A, Table S2**). Interestingly, we observed no significant enrichment of biological processes related to chemotaxis after 48 h, which could be an indication for a decreased cellular reaction towards the presence of the *L. casei* MVs. Additionally, we observed that genes, which were deregulated after transfection of *L. casei* MV-RNA for 24 h, were also involved in pathways like cell adhesion, and chemotaxis (**Figure 3F, Figure S4B-D, Table S2**) leading to the suggestion that *L. casei* MV-RNA independently causes transcriptional changes in similar pathways like MVs itself. For Caco-2 cells that had been incubated with *E. faecalis* MVs, we observed no enrichment in biological processes related to an altered immune response after 10 h, and only minor changes in genes related to inflammatory response or cell adhesion after 24 h incubation (**Figure 3G, Figure S4E, Table S2**). Remarkably, transfection of *E. faecalis* MV-RNA led to significant changes in the transcriptome after already 10 h, which included biological processes like defense response, cell adhesion, and chemotaxis (**Figure 3H, Figure S4F, Table S2**). Thus, it seems that the MV-RNA has a strong influence on transcriptomic changes in the Caco-2 cells. For cells that had been incubated with *P. mirabilis* OMVs for 10 h and 24 h, the differentially expressed genes were again significantly enriched in biological processes like inflammatory response, cell adhesion, and chemotaxis (**Figure 3I, Figure S4G, Table S2**). Similarly, deregulated genes after transfection of *P. mirabilis* OMV-RNA were again also involved in similar pathways indicating that the OMV-RNA might independently cause transcriptional changes like the OMVs itself (**Figure 3J, Figure S4H, Table S2**).

Given the presence of LPS in *P. mirabilis* OMVs and OMV-RNA, we additionally asked to which extent LPS contributes to the observed transcriptomic changes in Caco-2 cells. Our results revealed that the incubation or transfection of Caco-2 cells with *P. mirabilis* LPS caused deregulation of genes enriched in biological processes, which were rather dissimilar to those observed after treatment with *P. mirabilis* OMVs or OMV-RNA (**Figure S5**). We further investigated the expression of specific genes related to immune response after incubation or transfection of Caco-2 cells with *P. mirabilis* OMVs or OMV-RNA compared to LPS. We thereby computed a strong increase of the expression of i. e. *CCL20*, *CCL22*, *CXCL1*, *CXCL10*, *CXCL8*, *TLR3*, and *TLR7* after incubation of cells for either 10 h or 24 h with OMVs. After incubation with LPS from P*. mirabilis*, the expression of these genes remained mostly unchanged. Additionally, transfection of Caco-2 cells with OMV-RNA led to changes in the expression of genes like *CCR7*, *CX3CL1* or *IL-27*, compared to cells transfected with LPS from *P. mirabilis* (**Table S3**). In summary, our results indicate that *P. mirabilis* OMVs and OMV-RNA can cause transcriptional changes in the Caco-2 cells, which were independent from the effects caused by LPS only.

### Bacteria show interaction with Caco-2 derived EVs and altered bacterial growth

In order to facilitate cross-kingdom communication, bacteria must be able to interact with eukaryotic EVs. Thus, we asked whether bacteria take up Caco-2 derived EVs and if this influences their growth. To prevent influences by FCS-derived EVs, we used EVs from Caco-2 that were cultured in FCS-free medium. Isolated Caco-2 (-FCS) EVs had a size of 121.2 (± 6.1) nm and a concentration of 1.2x10^11^ (± 2.7x10^10^) particles/ml (**Figure 4A and 4B**).

**Figure 4:**
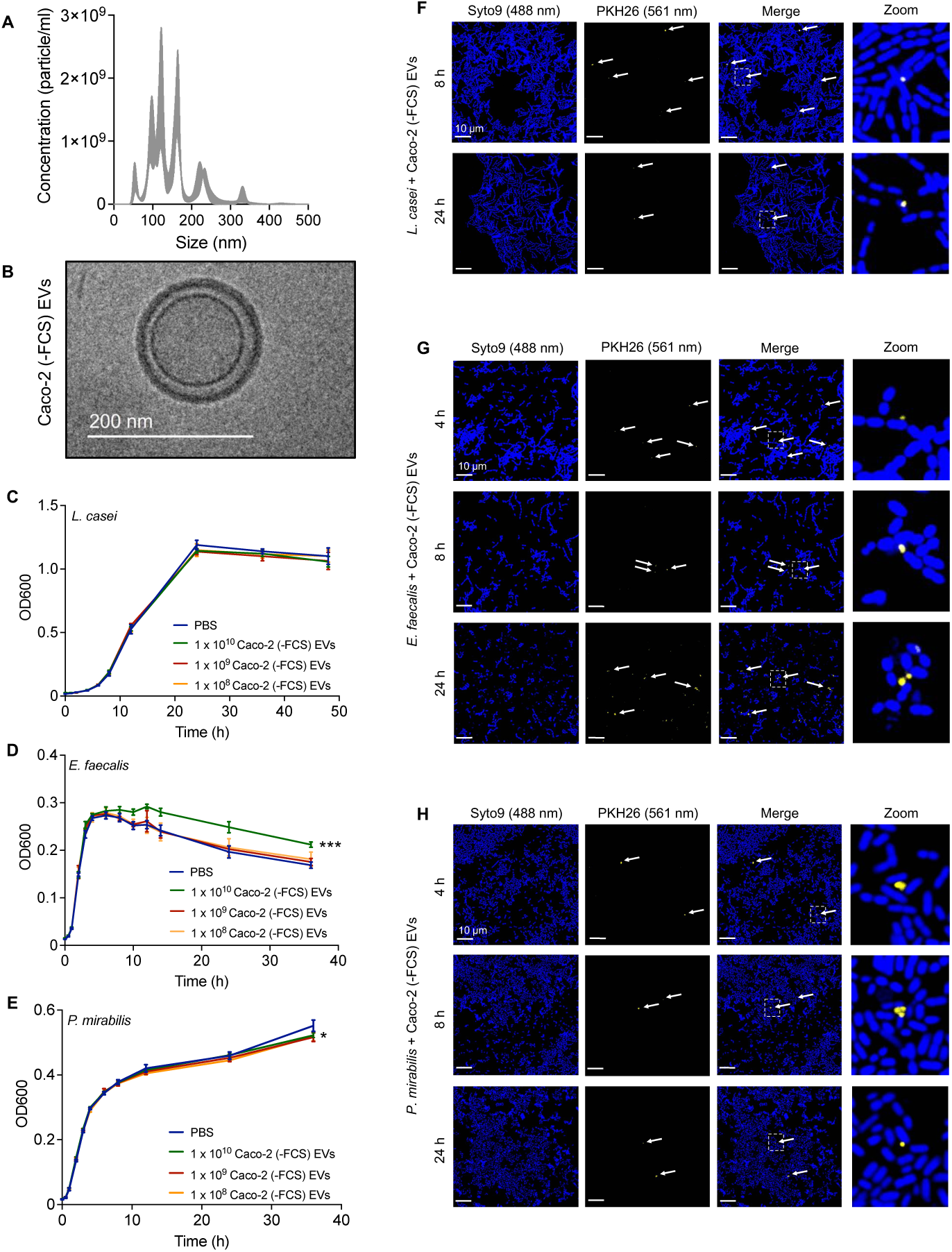
Uptake of Caco-2 EVs by *L. casei*, *E. faecalis* and *P. mirabilis*. (**A and B**) Characterization of Caco-2 EVs. (**A**) Typical size distribution of purified Caco-2 (-FCS) EVs measured by nanoparticle tracking analysis. (**B**) Representative cryo-TEM images of Caco-2 (-FCS) EVs showed round shaped particles. (**C-E**) Growth of *L. casei* (**C**), *E. faecalis* (**D**), and *P. mirabilis* (**E**) in the presence of different amounts of Caco-2 (-FCS) EVs or PBS as a control over a period of 48 h or 36 h at 37 °C. Results are presented as mean ± SD from 4 independent experiments. (**F-H**) Visualization of the interaction of *L. casei* (**F**), *E. faecalis* (**G**) or *P. mirabilis* (**H**) with Caco-2 (-FCS) EVs. Bacteria were incubated for 4 h, 8 h or 24 h with PKH26-labeled Caco-2 (-FCS) EVs. After the incubation, bacteria were stained with SYTO™ 9 and fixed. SYTO™ 9 was visualized using an argon laser at 488 nm, and PKH26 with a DPSS laser at 561 nm. For better illustration, the bacteria are shown in blue, and EVs are displayed in yellow. The white arrows indicate areas where PKH26-labeled Caco-2 (-FCS) EVs were detected. Scale bar: 10 µm. For all experiments, a two-tailed unpaired t-test was conducted to determine statistical significance (p < 0.05 *, p < 0.001 ***).

Remarkably, we observed that *E. faecalis* cultivated with 1 x 10^10^ Caco-2 EV exhibited a prolonged stationary phase before entering death phase. The growth of *L. casei* was not affected when cultivating the bacteria in the presence of different amounts of Caco-2 EVs. Likewise, we initially detected no effect of the EVs on the growth for *P. mirabilis*. It took 36 h until we observed a reduced growth of the bacteria cultivated with Caco-2 EVs (**Figure 4C-E**). To further investigate how the Caco-2 EVs might have affected the bacterial growth, we asked whether the bacteria are able to take up eukaryotic EVs. Therefore, we cultivated the bacteria with PKH26-labeled Caco-2 EVs. For *L. casei* and *P. mirabilis* we detected only a small amount of EVs next to or in proximity to the bacteria. However, for *E. faecalis*, we observed an increasing amount of EVs in the vicinity or co-localizing with the bacteria (**Figure 4F-H**). Concerning to the growth curve, the delayed entry of *E. faecalis* into the death phase might be due to the uptake of the Caco-2 EVs.

### EVs from conditioned and unconditioned media exhibit different miRNA expression profiles

Recent studies suggest that miRNAs secreted within eukaryotic EVs are functionally delivered to recipient cells [5]. A common approach to study the complex functions of EVs in cell-to-cell communication is to use EVs derived from cell culture media [25]. The major challenge regarding the analysis of EVs and especially EV-miRNAs from cell culture is the presence of FCS-derived EVs [26]. To date, numerous alternatives, i. e. use of serum-free media or ultracentrifugation of FCS before use, have been proposed to overcome this challenge [27]. However, the depletion of FCS can cause reduced growth and viability of the cells [28] as well as altered packaging of miRNAs in EVs [29]. We thus opted for a different approach to identify miRNAs that are secreted within EVs from Caco-2 cells, which avoids the depletion of FCS from the cell culture system. Therefore, we first isolated EVs from Caco-2 cells cultured with FCS-containing medium and additionally from FCS-containing DMEM itself. Caco-2 (+FCS) EVs showed a size of 123.1 (± 10.4) nm and a concentration of 1.1x10^11^ (± 2.8x10^10^) particles/ml. DMEM (+FCS) EVs had a similar size of 117.4 (± 6.9) nm and concentration of 7.4x10^10^ (± 1.3x10^10^) particles/ml (**Figure 5A-D**). By using western blot analysis, we confirmed the presence of EVs marker and absence of cellular marker (**Figure S6**). Using smallRNA-seq, we then sequenced miRNAs isolated from Caco-2 (+FCS) EVs (conditioned medium) and compared it to the miRNA expression profile of DMEM (+FCS) EVs (unconditioned medium).

**Figure 5:**
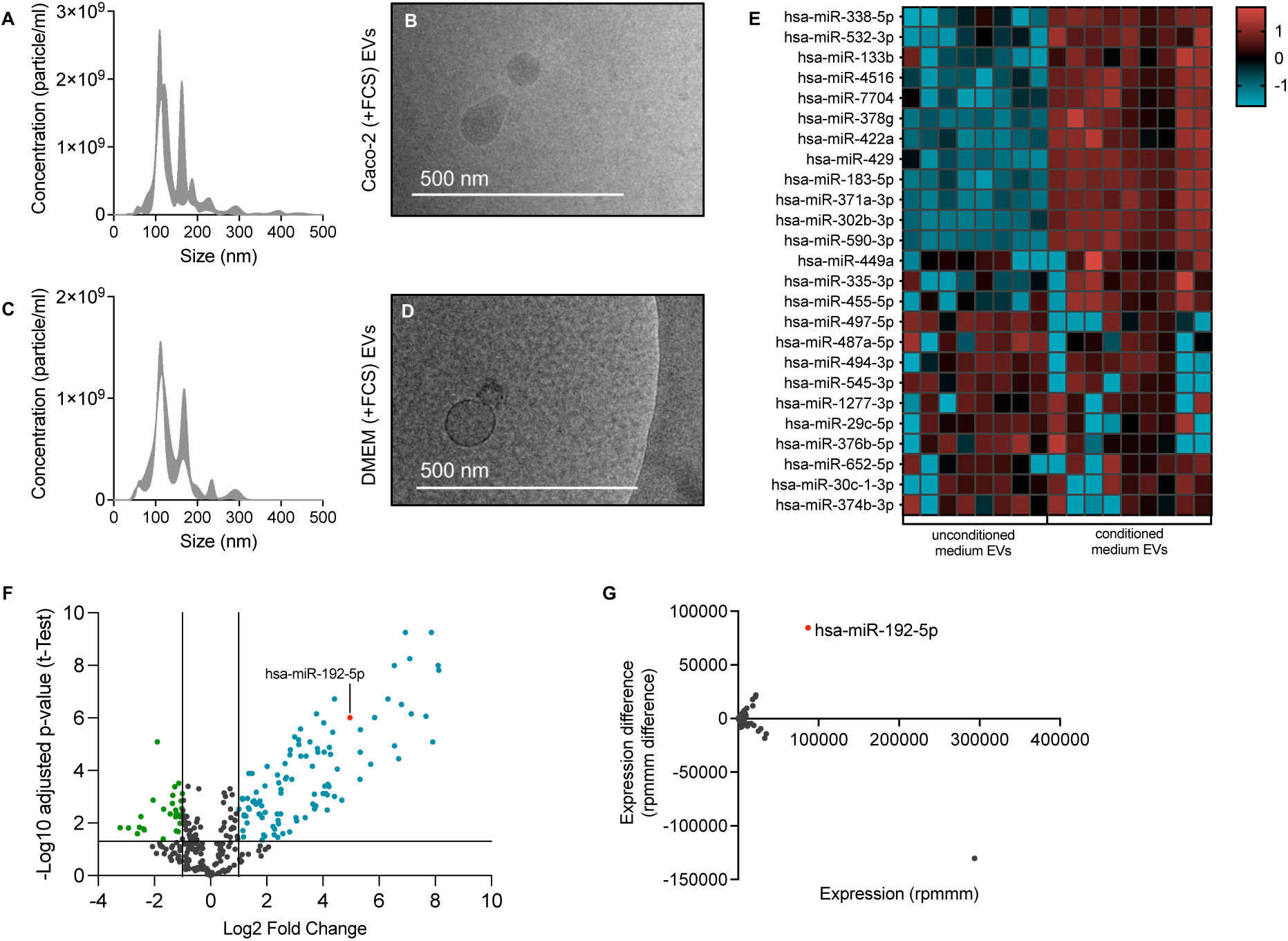
miRNA expression profiles of conditioned and unconditioned medium EVs. (**A-D**) Characterization of Caco-2 EVs and DMEM EVs. Typical size distribution of purified Caco-2 (+FCS) EVs (**A**) and DMEM (+FCS) EVs (**C**) measured by nanoparticle tracking analysis. Representative cryo-TEM images of Caco-2 (+FCS) EVs (**B**) and DMEM (+FCS) EVs (**D**) showed round shaped particles. (**E-G**) Analysis of smallRNA-seq data. (**E**) Heatmap of top 25 expressed miRNAs based on hierarchical clustering analyses. (**F**) Scatter plot for miRNAs detected in at least 75 % of EVs from conditioned and unconditioned medium. X-axis shows the Log2 fold change, and y-axis shows the -Log10 of the Benjamini-Hochberg-adjusted p-value. Each dot represents a single miRNA. Displayed in green are miRNAs whose expression is significantly (p < 0.05) and at least two-fold reduced (Log2 fold change < -1) in the EVs from the conditioned medium. Displayed in blue are miRNAs whose expression is significantly (p < 0.05) and at least two-fold increased (Log2 fold change > 1) in the EVs from the conditioned medium. MiR-192-5p is displayed in red. (**G**) Scatter plot for miRNAs detected in at least 1 sample of EVs from conditioned and unconditioned medium. X-axis shows the expression (rpmmm), and y-axis shows the expression difference (rpmmm difference). Each dot represents a single miRNA. MiR-192-5p is displayed in red. An expression difference < 0 indicates a reduced expression (rpmmm), an expression difference > 0 indicates an increased expression (rpmmm) of the corresponding miRNA in EVs from the conditioned medium.

Performing a hierarchical cluster analysis, we found a discernible clustering pattern based on whether EVs were isolated from conditioned or unconditioned medium (**Figure 5E**). Using differential expression analysis, we detected a total of 316 miRNAs in EVs from both conditions (**Table S4**). Moreover, we found that 104 miRNAs showed an at least 2-fold increased expression (Log2 FC > 1 and p < 0.05) in EVs from conditioned medium indicating an elevated probability that these miRNAs are released in EVs from the Caco-2 cells. Contrary, we found 32 miRNAs that showed a significantly reduced expression (Log2 FC < -1 and p < 0.05) in conditioned compared to unconditioned medium EVs making an origin from Caco-2 cells highly unlikely (**Figure 5F**). Towards further detection of miRNA that are most likely secreted in EVs by Caco-2 cells we determined how often a given miRNA was read in relation to 1,000,000 reads (rpmmm, reads per mapped million miRNAs). Among these miRNAs, we identified miR-192-5p, which was frequently detected in the EVs from the conditioned medium (**Figure 5G**).

### miR-192-5p is selectively taken up, and packaging in liposomes influences their uptake by bacteria

Since the analysis of the miRNA content in EVs revealed the elevated presence of miR-192-5p in EVs from Caco-2 cells, we proposed that this miRNA had the highest potential to affect the bacteria. To investigate whether miR-192-5p influences the growth of the bacteria, we cultivated them in the presence of different concentrations of synthetic miR-192-5p. While the growth of *L. casei* and *E. faecalis* remained stable, we detected that the presence of 4 µM synthetic miRNA significantly promoted the growth of *P. mirabilis* (**Figure 6A-C**). We then asked whether the promoted growth was due to an increased number of live bacteria. However, the percentage of live bacteria remained the same for all three tested bacteria after incubation with 4 µM synthetic miRNA. This led us to the suggestion that the increased optical density of *P. mirabilis* cultures was rather due to some other factors most likely not related to an increased percentage of viable bacteria. (**Figure S7A-C**).

**Figure 6:**
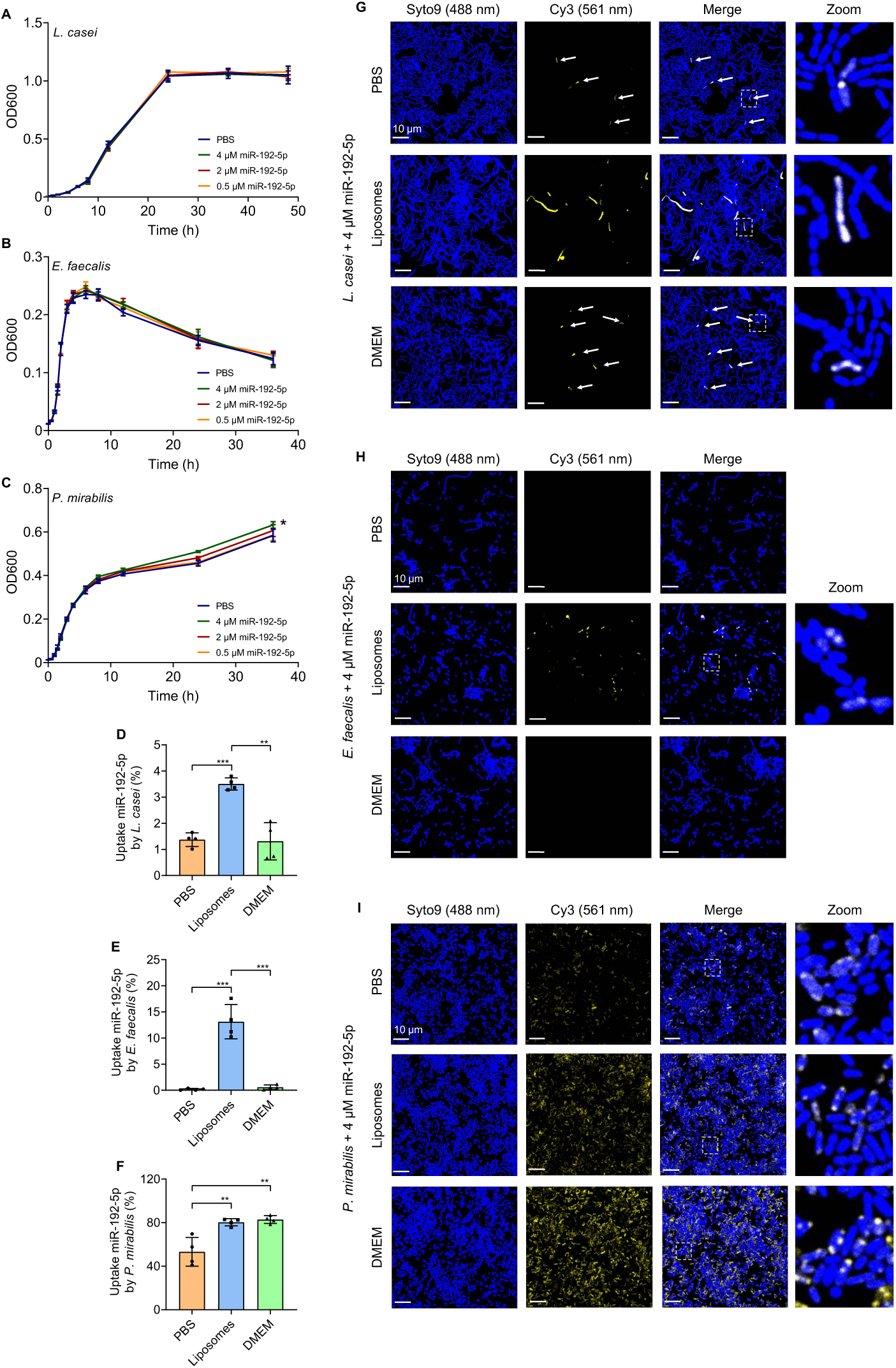
Uptake of free and liposome-packaged miR-192-5p by *L. casei*, *E. faecalis* and *P. mirabilis*. (**A-C**) Growth of *L. casei* (**A**), *E. faecalis* (**B**), and *P. mirabilis* (**C**) in the presence different concentrations of synthetic miR-192-5p or PBS as a control over a period of 48 h or 36 h at 37 °C. Results are presented as mean ± SD from 3-4 independent experiments. (**D-F**) Flow cytometry analysis of the uptake of fluorescence-labeled synthetic miRNA by various bacteria. All bacteria were cultivated in the presence of 4 μM free Cy3-labeled synthetic miR-192-5p in PBS or DMEM, or 4 μM liposome-packaged Cy3-labeled synthetic miR-192-5p for 24 h (*E. faecalis* or *P. mirabilis*) or 48 h (*L. casei*) at 37 °C. After the incubation, bacteria were stained with SYTO™ 9 and fixed. The percentage of SYTO™ 9/PE (Phycoerythrin)-positive bacteria compared to the respective control (PBS, DMEM or liposomes). Results are presented as mean ± SD from 4 independent experiments. (**G-I**) Visualization of the interaction of *L. casei* (**G**), *E. faecalis* (**H**) or *P. mirabilis* (**I**) with free or liposome-packaged fluorescently labeled synthetic miRNA. Bacteria were cultivated in the presence of 4 μM free Cy3-labeled synthetic miR-192-5p in PBS or DMEM, or 4 μM liposome-packaged Cy3-labeled synthetic miR-192-5p for 24 h (*E. faecalis* or *P. mirabilis*) or 48 h (*L. casei*) at 37 °C. After the incubation, bacteria were stained with SYTO™ 9 and fixed. SYTO™ 9 was visualized using an argon laser at 488 nm, and Cy3 with a DPSS laser at 561 nm. For better illustration, the bacteria are shown in blue, and synthetic miRNA is displayed in yellow. The white arrows indicate areas where weak signals for Cy3-labeled synthetic miR-192-5p were detected. Scale bar: 10 µm. For all experiments, a two-tailed unpaired t-test was conducted to determine statistical significance (p < 0.05 *, p < 0.01 **, p < 0.001 ***).

To examine whether the bacteria are still able to internalize the miRNA, we cultivated them in the presence of 4 µM Cy3-labeled synthetic miRNA. Flow cytometry analysis (**Figure 6D-F**) and CLSM (**Figure 6G-I**) showed that *L. casei* and *E. faecalis* hardly took up the synthetic miRNA whereas 53.1 (± 13.2) % of *P. mirabilis* showed uptake and co-localization with the synthetic miR-192-5p. Our finding indicates that the bacteria have different abilities to internalize the free miRNA.

Since we found that miR-192-5p is present in EVs released from Caco-2 cells, this promoted us to ask to which extent packaging of miRNA influences their uptake by bacteria. Thus, we cultivated the bacteria in the presence of 4 µM liposome-packaged synthetic miRNA to mimic EVs. First, we examined whether the growth of the bacteria might be affected. We found a mild decrease in the percentage of live *L. casei* after incubation with the empty liposomes, which we did not observe when the bacteria were incubated with synthetic miRNA-containing liposome. To determine if the increased number of viable bacteria was specifically due to the presence of miR-192-5p in the liposomes, we used polyadenylic acid (Poly(A)), which is also able to form a complex with the liposomes. We observed that Poly(A)-containing liposomes also reduced the proportion of live *L. casei* indicating that the synthetic miRNA-containing liposomes might have a positive effect on the number of viable bacteria. As for *E. faecalis*, the percentage of live bacteria remained stable after incubation with any of the liposomes. However, for *P. mirabilis* on the other hand, we observed an increase of viable bacteria after incubation with liposome-packaged synthetic miRNA, Poly(A)-containing liposomes, and empty liposomes (**Figure S7D-F**).

Additionally, we analyzed the internalization of the liposome-packaged miRNA after cultivation of the bacteria in the presence of 4 µM liposome-packaged synthetic miR-192-5p. Strikingly, we found that the liposomes increased the ability of bacteria to internalize the miRNA. We detected uptake of liposome-packaged miRNA in 3.5 (± 0.2) % of *L. casei* (**Figure 6D**), 13.1 (± 3.3) % of *E. faecalis* (**Figure 6E**) and 80.2 (± 3.4) % of *P. mirabilis* (**Figure 6F**). In accordance with these results, we observed an increased accumulation of liposome-packaged synthetic miR-192-5p in the bacteria using CLSM (**Figure 6G-I**)

To further elucidate whether the miRNA packing impacts the miRNA uptake, we determined the impact of the incubation media. For the experiments with the liposomes, the bacteria were cultivated in a mixture of their respective growth medium and DMEM, which was used for liposome-complex preparation. For the experiments with the free miRNA, however, the miRNA was added to the respective medium together with PBS. To exclude the possibility that DMEM had an influence on the ability of the bacteria to take up the miRNA, we examined the uptake of the free miRNA in DMEM. Remarkably, for *L. casei* and *E. faecalis* we detected a rather similar uptake level for the free miRNA in DMEM, which was comparable to bacteria that had been cultivated in the presence of free miRNA in PBS. Interestingly, for *P. mirabilis* we found that also 82.8 (± 3.6) % of the bacteria had taken up the free miRNA in DMEM (**Figure 6D-I**) indicating that DMEM may affect the uptake of the miR-192-5p by *P. mirabilis*. In summary, our findings suggest that packaging of miR-192-5p in EV-like particles increases the uptake by bacteria to a rather different extent for each of the tested bacteria with the most pronounced effects in *L. casei* and *E. faecalis*.

## Discussion

Cross-kingdom communication via EVs represents an emerging mechanism by which microbial and host cells exchange molecular information [2]. While previous studies have described EVs in cross-kingdom contexts, our work integrates these observations into a unified, bidirectional framework. We propose that EVs and their RNA cargo act not merely as passive transporters, but as directionally active messengers, selectively influencing gene expression and growth behavior in recipient cells across kingdom boundaries.

Our data show that BEVs from *E. faecalis* and *P. mirabilis* are rapidly internalized by Caco-2 cells, whereas MVs from *L. casei* are taken up much more slowly. Differences in uptake may be related to the presence of virulence factors like cell wall polysaccharides in the cell envelope of *E. faecalis*, which plays a major role in host-pathogen interaction [30] or LPS in the outer membrane of Gram-negative bacteria, like *P. mirabilis*, which are recognized by Toll-like receptor 4 (TLR4) [31]. Using RNA-seq, we reveal that BEVs induce distinct transcriptional changes in Caco-2 cells, particularly in genes related to immune response such as inflammatory response or defense response as well as chemotaxis. Remarkably, we find that BEV-RNA alone is able to cause similar effects, providing mechanistic evidence that RNA cargo contributes independently to the cellular response. These observations support previous studies reporting that BEV-RNA can alter gene expression in recipient cells [10–12]. Furthermore, we observe that the effects of *P. mirabilis* OMVs and OMV-RNA differ from those caused by *P. mirabilis* LPS alone, suggesting a distinct role of OMVs and their RNA content in regulating gene expression of host cells. However, another study has reported that LPS and OMV-RNA isolated from *Escherichia coli* OMVs can induce similar gene expression changes in bladder cells [32].

Conversely, we demonstrate that *E. faecalis* can internalize Caco-2-derived EVs and that cultivation of *E. faecalis* in the presence of these EVs promotes bacterial growth. SmallRNA-seq revealed that miR-192-5p, which is known to be highly expressed in colon [33], is frequently abundant in Caco-2 EVs. The uptake of mammalian miRNAs by bacteria has thus far been reported in only a few cases [16–18, 20]. In our study, we observe species-specific differences in miR-192-5p uptake, with Gram-negative bacterium *P. mirabilis* showing the highest uptake rates. CLSM reveals that miR-192-5p mostly accumulates at the poles of *P. mirabilis*. Given that pili are often localized at bacterial poles [34, 35], uptake may be mediated by type IV pili, which can be used by some bacteria to take up extracellular DNA [36]. The cell wall of Gram-positive bacteria such as *L. casei* and *E. faecalis* consists of a thick peptidoglycan layer, as well as teichoic acids, wall polysaccharides, and proteins [37]. Additionally, *E. faecalis* possesses several DNA defense mechanisms, making the uptake of foreign nucleic acids even more difficult [38]. Notably, we find that packaging of miR-192-5p in artificial liposomes significantly enhance its uptake by *L. casei* and *E. faecalis*. This observation aligns with recent finding showing that lipid-polymer hybrid nanoparticles could improve the delivery of ampicillin to *E. faecalis*, whereas the delivery failed when the bacteria were incubated with free ampicillin [39]. These results suggest that Gram-positive bacteria are more likely to internalize miRNAs when encapsulated within EVs, rather than as free molecules circulating in the gut lumen.

Taken together, this study reveals EVs as bidirectional molecular messengers in host-microbe communication network. These insights not only expand our understanding of host-microbiome interactions but also lay the groundwork for exploring EVs as programmable carriers in targeted therapies. Future studies should investigate whether these interactions occur *in vivo*, in the presence of immune components and under disease-relevant conditions. Nevertheless, our work introduces a conceptual model in which EVs act as mediators in the molecular dialogue between humans and their microbiota.

## Methods

### Bacterial Cultures

*L. casei* (DSM 20011) was grown in deMan, Rogosa and Sharpe (MRS) broth (Carl Roth), *E. faecalis* (DSM 20478) and *P. mirabilis* (DSM 4479) were grown in brain heart infusion (BHI) broth (Merck Millipore). Bacterial growth was determined by measuring the optical density (OD) at a wavelength of λ = 600 nm or the colony forming units per ml (CFU/ml). Liquid cultures were cultivated under static conditions at 37 °C starting with an OD of 0.1 until they reached the stationary phase of bacterial growth.

### Eukaryotic Cell Culture

Human colon adenocarcinoma cells, Caco-2 (ATCC HTB-37), were maintained in DMEM (Thermo Fisher Scientific) supplemented with 10 % (v/v) FCS (Thermo Fisher Scientific) and 1 % (v/v) non-essential amino acids (Thermo Fisher Scientific) at 37 °C and 5 % CO_2_. Medium was exchanged every 2-3 days, and the cells were split once a week. All experiments were performed during passage 19 ± 5.

### Isolation of proteins from eukaryotic cells

Caco-2 cells were grown in cell culture medium. Medium was exchanged every 2-3 days. After 7 days, the medium was aspirated, cells were washed twice with PBS and detached using Trypsin-EDTA (Thermo Fisher Scientific). Cells were further pelleted at 300 x g for 4 min, resuspended in fresh cell culture medium and counted using the CASY Cell counter & Analyzer (OLS OMNI Life Science). A total of 6 x 10^6^ cells were transferred into a reaction tube and washed with PBS twice. Next, cells were resuspended in 100 µl RIPA buffer (Thermo Fisher Scientific) supplemented with cOmplete™ protease inhibitor cocktail (Roche), sonicated at 10 % for 10 x 2 s at intervals of 10 s and incubated in the fridge at 4 °C overnight. On the following day, samples were centrifuged at 12.000 x g and 4 °C for 10 min. Supernatant was transferred into a new reaction tube and stored at -80 °C.

### BEV Isolation and Purification

Bacterial cultures were centrifuged to pellet the bacteria. Supernatants were sterile filtered through a 0.22 µm (*P. mirabilis*) or 0.45 µm (*L. casei* and *E. faecalis*) PDVF membrane (Merck Millipore) to remove remaining bacteria. Filtered supernatants were transferred into 70-ml polycarbonate ultracentrifuge tubes (Beckman Coulter) and centrifuged at 100.000 x g at 4 °C for 2 hours (*L. casei* and *P. mirabilis*) or 160.000 x g at 4 °C for 3 hours (*E. faecalis*). After ultracentrifugation, the supernatant was discarded and the pellets from 3 ultracentrifuge tubes were pooled and resuspended in a total of 500 µl filtered PBS (Thermo Fisher Scientific). The resuspended pellets were either stored at -80 °C or directly purified under aseptic conditions by an SEC column filled with 40 ml Sepharose CL-2B (Sigma-Aldrich) in PBS. Fractions of 1 ml each were collected and stored at -80 °C until further use.

### EV Isolation and Purification

EVs were isolated from three different conditions. Cell culture supernatant from cells grown in cell culture medium with supplements (Caco-2 (+FCS), conditioned medium) was collected after around 72 h of growth on the day of cell passaging (day 7). Cells were pelleted at 300 x g for 4 min and cell-free supernatant was stored at -80 °C until further processing. For isolation of EVs from cells grown without supplements (Caco-2 (-FCS)), cells were maintained in cell culture medium with supplements for 7 days. Then, the cells were washed twice with PBS and cell culture medium without supplements was added to the cells. After 24 h at 37 °C and 5 % CO_2_, supernatant was collected, cell culture medium without supplements was added to the cells and the supernatant was again collected after 24 h. As described above, all supernatants were centrifuged at 300 x g for 4 min to pellet the cells and cell-free supernatant was stored at -80 °C. For EV isolation, cell-free supernatants were thawed at RT and centrifuged at 3.000 x g for 20 min at 4 °C to pellet larger particles. In parallel, fresh cell culture medium supplemented with FCS (DMEM (+FCS), unconditioned medium) was also centrifuged at 3.000 x g for 20 min at 4 °C to ensure similar initial preparation of both media. The following steps were carried out in the same manner for all EV preparations. Next, the supernatant was carefully transferred into 70-ml polycarbonate ultracentrifuge tubes (Beckman Coulter) and centrifuged at 100.000 x g at 4 °C for 2 hours. After ultracentrifugation, the supernatant was removed and the EV pellets from 3 ultracentrifuge tubes were pooled and resuspended in a total of 500 µl filtered PBS (Thermo Fisher Scientific). Only the pelleted Caco-2 (-FCS) EVs that were used for visualizing the uptake of EVs into bacteria were resuspended in 200 µl filtered PBS. The resuspended pellets were either stored at -80 °C or directly purified using an SEC column filled with 40 ml Sepharose CL-2B (Sigma-Aldrich) in PBS. Fractions of 1 ml each were collected and stored at -80 °C until further use.

### Determination of Protein Content

The protein concentration of the different EV and BEV fractions after SEC and Caco-2 protein extract was determined using QuantiPro BCA Assay Kit (Sigma-Aldrich) according to manufacturer’s recommendations.

### Endotoxin Quantification

Endotoxin (Lipopolysaccharide, LPS) concentration of *P. mirabilis* OMV and *P. mirabilis* OMV-RNA was determined using Pierce™ Chromogenic Endotoxin Quant Kit (Thermo Scientific). In brief, 50 µl of each OMV sample and 1 µl of each OMV-RNA sample was used, and the assay was performed according to manufacturer’s instructions. Endotoxin concentration was expressed in endotoxin units (EU) per ml and transformed in nanogram per ml using the equation 10 EU/ml equals 1 ng/ml.

### Determination of Particle Concentration and Size

The concentrations and sizes of the EVs and BEVs were determined using NanoSight (Malvern Panalytical). Before measurement, the samples were diluted with filtered PBS, and 400 µl of diluted sample was introduced into the measuring chamber. Videos with a length of 30 sec were recorded in triplicates with a camera level of 15. Particle concentrations and sizes were calculated by NTA 3.4 software version 3.4.4 with a detection threshold set to 5.

### Cryo-Transmission Electron Microscopy (Cryo-TEM)

For cryo-TEM a 2 µl droplet of the sample was placed on a holey carbon supported copper grid (Plano, type S147-4), blotted for 2 s and vitrified in undercooled liquid ethane using a Gatan CP3 plunge-freezer. The frozen hydrated specimen was transferred to a Gatan model 914 cryo-TEM sample holder under liquid nitrogen and visualized using a JEOL JEM-2100 LaB6 TEM at 200 kV accelerating voltage under low-dose conditions (CCD camera Gatan Orius SC1000, 2 s acquisition time).

### Scanning Electron Microscopy (SEM)

For SEM, overnight cultures from all three bacteria were diluted with fresh broth to 0.1 in a total volume of 2 ml and cultivated at 37 °C. After 12 h (*E. faecalis* and *P. mirabilis*) or 40 h (*L. casei*) the bacteria were fixed with 3 % (v/v) glutardialdehyde (Thermo Fisher Scientific) for 2 h at RT. Next, the samples were washed with increasing concentrations of ethanol up to 100 %, and hexamethyldisilazane (HMDS) (Sigma-Aldrich) was added for the final drying process. The HDMS was removed, and the samples were dried at RT for several hours. Afterwards, the samples were placed on sticky carbon tapes, and sputter coated with gold using Q150R ES (Quorum Technologies Ltd.). Images were taken with Evo HD 15 (Carl Zeiss Microscopy GmbH) and processed using SmartSEM software version 6.09.

### Western Blot

The presence of EV and cellular marker was determined by Western blot analysis. For gel electrophoresis, 15 µg Caco-2 protein extract and 30 µl of each EV sample was mixed with 2x Sample Buffer (130 mM Tris, 6 % (v/v) SDS, 10 % (v/v) 3-mercapto-1,2-propanediol, 10 % (v/v) glycerol). Samples were sonicated 3 x 3 s (40 %), incubated on ice for 5 min and at 99 °C for 10 min. Next, samples were centrifuged at 14.000 rpm and 4 °C for 10 min and supernatant was used for further analysis. Samples were separated using (4–15%) mini-PROTEAN® TGX™ Precast Protein (Bio-Rad) for 30 min at 12 mA and 40 min at 25 mA. Separated proteins were transferred onto a nitrocellulose membrane (GE Healthcare) for 2 h at 400 mA. Anti-Calnexin antibody (#2679, RRID:AB_2228381, Cell Signaling Technology) and anti-CD81 antibody (#56039, RRID:AB_2924772, Cell Signaling Technology) were diluted in 5 % (w/v) BSA and 1x TBST (1:800) and used to detect the protein level in Caco-2 protein extract and EV samples. A secondary anti-rabbit antibody was purchased from Sigma-Aldrich (A0545, RRID:AB_257896). Signals were captured using ChemiDoc™ Touch system (Bio-Rad).

### BEV-RNA Isolation and Quality Control

Total BEV-RNA was isolated from the 400 µl BEV fraction with the highest particle concentration using miRNeasy Serum/Plasma Kit (Qiagen) according to the manufacturer’s instructions with modifications and additional On-Column DNase digestion using RNA-free DNase Set (Qiagen). BEV-RNA concentration was measured using Qubit 1 x RNA HS Assay Kit (Thermo Fisher Scientific). BEV-RNA fragments were analyzed using the Agilent RNA 6000 Pico Kit for Agilent 2100 Bioanalyzer (Agilent Technologies).

### EV-RNA Isolation and Quality Control

Total EV-RNA was isolated from 400 µl EV fraction with the highest particle concentration using miRNeasy Serum/Plasma Kit (Qiagen) according to the manufacturer’s instructions with modifications. EV-RNA concentration was measured using Nanodrop 2000 spectrophotometer (Thermo Fisher Scientific). EV-RNA fragments were analyzed using the Agilent RNA 6000 Pico Kit for Agilent 2100 Bioanalyzer (Agilent Technologies).

### Eukaryotic Cell Viability and Cytotoxicity Assay

Cell viability and cytotoxicity was tested after incubation with BEVs or after transfection with BEV-RNA. Caco-2 cells were seeded with a density of 7.5x10^4^ cells/well into a 96-well plate and incubated at 37 °C and 5 % CO_2_. After 24 h, the medium was aspirated and 100 µl fresh medium without FCS was added to the cells together with 100 µl of the BEV sample or three serial 1:10 dilutions of the BEVs. Every experiment included a live-control using cells incubated with filtered PBS, and a dead-control using cells treated with 1 % (v/v) Triton X-100 (Sigma-Aldrich). To test the cell viability after transfection with BEV-RNA, a range of different amounts of BEV-RNA or *P. mirabilis* LPS (Sigma-Aldrich) were mixed with transfection reagent Lipofectamine™ 3000 (Thermo Fisher Scientific) according to manufacturer’s instructions and added to the cells along with fresh medium without FCS to a final volume of 200 µl. Again, every experiment included a live-control using cells incubated with transfection reagent mixed with nuclease-free water (vehicle), and a dead-control using cells treated with 1 % (v/v) Triton X-100 (Sigma-Aldrich).

After additional 24 h or 48 h, cytotoxicity was assessed using the Cytotoxicity Detection Kit (Roche). In brief, 100 µl of supernatant was mixed with 100 µl LDH working solution. After incubation at RT for 5 min under mild shaking the absorbance was measured at a wavelength of λ = 490 nm. To further test the viability of the cells, PrestoBlue reagent (Thermo Fisher Scientific) was diluted 1:10 with fresh medium. The remaining medium in the 96-well plate was aspirated and 100 µl of diluted PrestoBlue reagent was added carefully to the cells. After incubation for 20 min at 37 °C, the fluorescence (λEx/ λEm 560/590 nm) was measured.

### Flow Cytometry and Confocal Laser Scanning Microscopy (CLSM) of eukaryotic cells

Unpurified BEV pellets were incubated with 2 µl (*L. casei*) or 1 µl (*E. faecalis* and *P. mirabilis*) Vybrant™ DiI (Invitrogen) for 30 min at 37 °C. Stained BEVs were purified by an SEC column filled with 40 ml Sepharose CL-2B (Sigma-Aldrich) in PBS to remove free proteins and unbound dye. 50 µl of each collected fraction was used to measure the fluorescence (λEx/λEm 490/570 nm). The fraction with the highest fluorescence was used for further experiments.

For flow cytometry, cells were seeded in 48-well plates with 2x10^5^ cells/well and incubated at 37 °C and 5 % CO_2_. After 24 h, the cell culture medium was aspirated. Stained BEVs or PBS as a control were diluted 1:5 with fresh cell culture medium and added to the cells. After 4 h, 12 h, 24 h or 48 h the supernatant was removed, the cells were washed with PBS and detached using Trypsin-EDTA (Thermo Fisher Scientific). The cells were fixed with 4 % (v/v) formaldehyde (Thermo Fisher Scientific) for 30 min at RT, washed with PBS once and resuspended in 600 µl PBS. To detect DiI, a laser at 561 nm (PE, Phycoerythrin) was used (BD LSRFortessa™ Cell Analyzer). A total of 10.000 events per sample was set to be analyzed by BD FACSDiva™ software version 9.0.1 and evaluated by FlowJo™ software version 10.6.0.

For CLSM, the cells were seeded in 8-well chamber slides (Thermo Fisher Scientific) with a density of 2x10^5^ cells/well and incubated at 37 °C and 5 % CO_2_. After 24 h, the cell culture medium was aspirated and stained BEVs or PBS as a control were diluted 1:5 with fresh cell culture medium and added to the cells. After 12 h, 24 h or 48 h the cells were washed with PBS and fixed with 4 % (v/v) formaldehyde (Thermo Fisher Scientific) for 20 min at RT. Next, cells were permeabilized and unspecific binding sites were blocked using 1 % (w/v) BSA/0,05 % (w/v) saponin for 20 min at RT. F-actin was stained using Alexa Fluor™ 488 Phalloidin (Invitrogen) for 20 min and cell nuclei were stained using 4′, 6-Diamidino-2-phenylindole dihydrochloride (DAPI) (Sigma-Aldrich) for 20 min at RT. The cells were washed with PBS between the different staining steps. For confocal imaging (Leica TCS SP8 System) a laser at 405 nm was used to visualize DAPI, a laser at 488 nm for Phalloidin and a laser at 561 nm for DiI. The images were captured and processed using Leica Application Suite X software.

### Transcriptome analysis of eukaryotic cells

Caco-2 cells were seeded in 48-well plates with a density of 1.2x10^5^ cells/well and incubated at 37 °C and 5 % CO2. After 24 hours, the medium was aspirated and BEVs or BEV-RNA mixed with transfections reagent Lipofectamine™ 3000 (Thermo Fisher Scientific) were added together with fresh medium to the cells as described above. Controls used included PBS, *P. mirabilis* LPS (Sigma-Aldrich), and transfection reagent mixed with different amounts of AllStars Negative Control siRNA (ANC) (Qiagen) or *P. mirabilis* LPS. After 10 h, 24 h or 48 h, the supernatant was aspirated, and the cells were washed once with PBS. Subsequently, the cells were lysed by adding 700 µl Qiazol Lysis Reagent (Qiagen) to each well and stored at - 80 °C until RNA isolation. Total RNA was isolated from the Caco-2 cells using the miRNeasy Micro Kit (Qiagen) according to the manufacturer’s instructions. RNA concentration was measured by Nanodrop 2000 spectrophotometer (Thermo Fisher Scientific) and RNA integrity of random samples was tested using Agilent RNA 6000 Nano Kit (Agilent Technologies) on Agilent 2100 Bioanalyzer.

DNA libraries were prepared with the high-throughput MGISP-960 sample preparation system using the MGIEasy rRNA Depletion Kit (MGI Technologies) and MGIEasy RNA Library Prep Set (MGI Technologies) according to the manufacturer’s recommendations. Concentration of the PCR products was determined using Qubit 1 x dsDNA HS Assay Kit (Thermo Fisher Scientific). After PCR, different barcoded samples were pooled in equal amount and circularized to generate single stranded DNA libraries (ssDNA). The concentration of the ssDNA libraries was measured using Qubit ssDNA Assay Kit (Thermo Fisher Scientific). DNA nanoball (DNB) were generated by rolling circle amplification. Paired-end sequencing (PE100) was performed using DNBSEQ-G400RS High-throughput Sequencing Reagent Set (MGI Technologies) on the DNBSEQ-G400 instrument.

### EV miRNome analysis

DNA libraries were prepared using the MGIEasy smallRNA Library Prep Kit (MGI Technologies) according to the manufacturer’s recommendations. In brief, 100 ng of RNA from conditioned (Caco-2 (+FCS)) or unconditioned (DMEM (+FCS)) medium EVs was used as input. Amplified PCR products were separated using 6 % TBE gel electrophoresis. The target band from 100 to 120 bp was cut out and PCR products were eluted from the gel and purified by ethanol precipitation. Concentration of the PCR products was determined using Qubit 1 x dsDNA HS Assay Kit (Thermo Fisher Scientific). Next, different barcoded samples were pooled in equal amount and circularized to generate single stranded DNA libraries (ssDNA). Concentration of the ssDNA libraries was measured using Qubit ssDNA Assay Kit (Thermo Fisher Scientific). DNA nanoball (DNB) preparation and single-end sequencing (SE50) was carried out as a service at BGI (Shenzhen, China).

### Measurement of Bacterial Growth and Viability

For analysis of bacterial growth after incubation with EVs, overnight cultures of all three bacteria were prepared. On the following day, the OD was measured and cultures were diluted with fresh broth to 0.01. Next, EVs, two serial 1:10 dilutions of the EVs or PBS as a control were diluted 1:1 with bacterial suspension to a total volume of 200 µl. Bacteria were cultivated at 37 °C, and growth was determined by measuring the OD for 36 h or 48 h.

For analysis of bacterial growth and viability after incubation with synthetic miR-192-5p (Eurofins Genomics), overnight cultures of all three bacteria were prepared. On the following day, the OD was measured and cultures were diluted with fresh broth to 0.01. Next, 0.5 µM, 2 µM and 4 µM of synthetic miR-192-5p, or PBS as a control was added to a total volume of 150 µl. Bacteria were cultivated at 37 °C and growth was determined by measuring the OD for 36 h or 48 h. Bacterial viability was analyzed after 24 h (*E. faecalis* and *P. mirabilis*) or 40 h (*L. casei*) of growth in the presence of 4 µM synthetic miR-192-5p using Live/Dead™ BacLight™ Bacterial Viability Kit (Invitrogen) according to manufacturer’s recommendations. In brief, bacteria were stained with SYTO™ 9/PI for 15 min at RT and fluorescence of SYTO™ 9 (λEx/λEm 485/530 nm) and PI (λEx/λEm 485/630 nm) was measured. The percentage of live bacteria was determined using the adjusted dye ratio equation by Ou et al. [40].

For analysis of bacterial viability after incubation with liposome-packaged synthetic miR-192-5p, overnight cultures of all three bacteria were prepared. The OD was measured and cultures were diluted with fresh broth to 0.01. Next, 4 µM of synthetic miR-192-5p was mixed with Lipofectamine™ 3000 (Thermo Fisher Scientific) and was added to a total volume of 150 µl. Bacteria incubated with DMEM or Lipofectamine mixed with either 1x siRNA Max Buffer (supplied by Eurofins Genomics to resuspend the synthetic miR-192-5p) or 8 µg polyadenylic acid (Sigma-Aldrich) were used as controls. Bacterial viability was analyzed after 24 h (*E. faecalis* and *P. mirabilis*) or 40 h (*L. casei*) of growth using Live/Dead™ BacLight™ Bacterial Viability Kit (Invitrogen) according to manufacturer’s recommendations as described above.

### Flow Cytometry and Confocal Laser Scanning Microscopy (CLSM) of bacteria

For analysis of EV uptake by bacteria, EVs were stained using PKH26 (Sigma-Aldrich). According to manufacturer’s recommendations, PHK26 was diluted 1:125 (v/v) in Diluent C. Next, 200 µl unpurified Caco-2 (-FCS) EV pellet was incubated with 200 µl diluted PHK26 for 10 min at RT. Stained EVs were purified by an SEC column filled with 40 ml Sepharose CL-2B (Sigma-Aldrich) in PBS to remove free proteins and unbound dye. 50 µl of each collected fraction was used to measure the fluorescence (λEx/ λEm 490/570 nm). The fraction with the highest fluorescence was used for further experiments. The OD of overnight culture from all three bacteria was measured and cultures were diluted with fresh broth to 0.01. Next, stained EVs or PBS as a control were diluted 1:1 with bacterial suspension to a total volume of 200 µl and incubated at 37 °C. After 4 h, 8 h or 24 h, bacteria were stained with SYTO™ 9 for 10 min at 37 °C, fixed with 4 % (v/v) formaldehyde (Thermo Fisher Scientific) at 37 °C, washed once and then resuspended in PBS. 5 µl of each sample was applied on a coverslip together with a drop of fluorescence mounting medium (Agilent Technologies). For confocal imaging (Leica TCS SP8 System) a laser at 488 nm was used to visualize SYTO™ 9 and a laser at 561 nm for PKH26. The images were captured and processed using Leica Application Suite X software.

For analysis of miRNA uptake by bacteria, Cy3-labeled synthetic miR-192-5p (Eurofins Genomics) was used. The OD of overnight culture from all three bacteria was measured and cultures were diluted with fresh broth to 0.01. 4 µM of synthetic miR-192-5p in PBS, DMEM or mixed with Lipofectamine™ 3000 (Thermo Fisher Scientific) was added to a total volume of 150 µl. Bacteria incubated with PBS, DMEM or Lipofectamine mixed with 1x siRNA Max Buffer were used as controls. After 4 h, 8 h, 24 h or 48 h, bacteria were stained with SYTO™ 9 for 10 min at 37 °C, fixed with 4 % (v/v) formaldehyde (Thermo Fisher Scientific) at 37 °C, washed once and then resuspended in PBS. 5 µl of each sample was applied on a coverslip together with a drop of fluorescence mounting medium (Agilent Technologies). For confocal imaging (Leica TCS SP8 System) a laser at 488 nm was used to visualize SYTO™ 9 and a laser at 561 nm for Cy3. The images were captured and processed using Leica Application Suite X software. The residual sample was used for flow cytometry. To detect Cy3, a laser at 561 nm (PE, Phycoerythrin), and to detect SYTO™ 9, a laser at 488 nm (Alexa Fluor 488) was used (BD LSRFortessa™ Cell Analyzer). A total of 100.000 events (*E. faecalis*) or 200.000 events (*L. casei* or *P. mirabilis*) per sample was set to be analyzed by BD FACSDiva™ software version 9.0.1 and evaluated by FlowJo™ software version 10.6.0.

### Data analysis

Snakepipes pipeline (v2.7.3) [41] was used to process the paired-end Fastq files. Multiqc (v1.12) [42] was used to carry out initial QC. STAR (v2.7.10b) [43] was used to align the reads against GRCh38 using default options. Deduplication was performed with the markdup function from sambamba (v.0.8.0) [44]. FeatureCount from subread (v.2.0.1) [45] was used to quantify mRNA counts with the parameters -C -Q 10 --primary. The raw counts were used to perform differential expression analysis using DESeq2 (v1.30.1) [46]. Alongside the fold-change values provided by DESeq2 we calculate Cohen’s D using the variance stabilizing transform normalized counts created by Deseq2. We picked deregulated genes based on an absolute Log2 FC > 1 and a Cohen’s *d* > 0.8. The resulting lists were used to calculate gene set overlaps. GeneTrail 3.2 [24] was used for pathway analysis of differentially expressed genes by performing an over-representation analysis (ORA) with the following setting (Null hypothesis (for p-value computation): Two-sided; Method to adjust p-values: Benjamini-Hochberg; Significance level: 1; Minimal size of category: 1; reference set: all supported genes).

For miRNAs, miRmaster 2.0 pipeline [47] was used to trim, collapse and quantify the single-end sequencing fastq files using miRBase 22.1 [48]. The counts were normalized using reads per million mapped miRNA (rpmmm). MiRNA were filtered based on whether they were expressed in 75% of the groups. Differential expressions were calculated using t-test, followed by a Benjamini-Hochberg correction with an FDR of 10%.

Snakemake (v8.2.3) [49], Python (v3.12.2), matplotlib (v3.9.1) [50], seaborn (v0.13.2) [51], upsetplot (v0.9.0), R (v4.0.2), ggplot2 (v3.3.6) [52] were used for calculations and plotting.

## Resource availability

### Lead contacts

Requests for further information and resources should be directed to and will be fulfilled by the lead contacts, Laura Gröger, Andreas Keller, or Eckart Meese.

### Data and code availability

- RNA-seq data have been deposited at GEO and are publicly available as of the date of publication.
- smallRNA-seq data have been deposited at GEO and are publicly available as of the date of publication.
- Original western blot images have been deposited at Mendeley Data and are publicly available as of the date of publication.
- Any additional information required to reanalyze the data reported in the paper is available from the lead contact upon request.

## Acknowledgements

The authors acknowledge HPC resources support with hardware funded by the DFG within project 469073465. This study was funded by the Hedwig-Stalter Foundation (2022) for **L.G.**. This work was supported by the UdS-HIPS TANDEM Initiative.

## Author contributions

**Laura Gröger**: Conceptualization, Methodology, Validation, Formal analysis, Investigation, Writing - Original Draft, Writing - Review & Editing, Visualization and Project administration. **Shusruto Rishik**: Software, Formal analysis, Data Curation, Writing - Review & Editing and Visualization. **Nicole Ludwig**: Conceptualization, Methodology, Investigation and Writing - Review & Editing. **Amila Beganovic**: Investigation and Writing - Review & Editing. **Marcus Koch**: Investigation and Writing - Review & Editing. **Stefanie Rheinheimer**: Investigation and Writing - Review & Editing. **Martin Hart**: Methodology, Investigation and Writing - Review & Editing. **Petra König**: Investigation and Writing - Review & Editing. **Tabea Trampert**: Investigation and Writing - Review & Editing. **Pascal Paul**: Investigation and Writing - Review & Editing. **Annette Boese**: Investigation and Writing - Review & Editing. **Claus-Michael Lehr**: Resources and Writing - Review & Editing. **Sören L. Becker**: Resources and Writing - Review & Editing. **Gregor Fuhrmann**: Conceptualization, Resources, Writing - Review & Editing, Supervision, Project administration and Funding acquisition. **Andreas Keller**: Conceptualization, Resources, Writing - Original Draft, Writing - Review & Editing, Supervision, Project administration and Funding acquisition. **Eckart Meese**: Conceptualization, Resources, Writing - Original Draft, Writing - Review & Editing, Supervision, Project administration and Funding acquisition.

## References

1. Armingol, E., et al., Deciphering cell-cell interactions and communication from gene expression. Nat Rev Genet, 2021. 22(2): p. 71–88.

2. Munhoz da Rocha, I.F., et al., Cross-Kingdom Extracellular Vesicles EV-RNA Communication as a Mechanism for Host-Pathogen Interaction. Front Cell Infect Microbiol, 2020. 10: p. 593160.

3. Yanez-Mo, M., et al., Biological properties of extracellular vesicles and their physiological functions. J Extracell Vesicles, 2015. 4: p. 27066.

4. Bartel, D.P., MicroRNAs: genomics, biogenesis, mechanism, and function. Cell, 2004. 116(2): p. 281–97.

5. O’Brien, K., et al., RNA delivery by extracellular vesicles in mammalian cells and its applications. Nat Rev Mol Cell Biol, 2020. 21(10): p. 585–606.

6. Bauman, S.J. and M.J. Kuehn, Purification of outer membrane vesicles from Pseudomonas aeruginosa and their activation of an IL-8 response. Microbes Infect, 2006. 8(9-10): p. 2400–8.

7. Canas, M.A., et al., Outer Membrane Vesicles From Probiotic and Commensal Escherichia coli Activate NOD1-Mediated Immune Responses in Intestinal Epithelial Cells. Front Microbiol, 2018. 9: p. 498.

8. Karthikeyan, R., et al., Transcriptome responses of intestinal epithelial cells induced by membrane vesicles of Listeria monocytogenes. Curr Res Microb Sci, 2023. 4: p. 100185.

9. Shen, Y., et al., Outer membrane vesicles of a human commensal mediate immune regulation and disease protection. Cell Host Microbe, 2012. 12(4): p. 509–20.

10. Choi, J.W., et al., Secretable Small RNAs via Outer Membrane Vesicles in Periodontal Pathogens. J Dent Res, 2017. 96(4): p. 458–466.

11. Koeppen, K., et al., A Novel Mechanism of Host-Pathogen Interaction through sRNA in Bacterial Outer Membrane Vesicles. PLoS Pathog, 2016. 12(6): p. e1005672.

12. Zhang, H., et al., sncRNAs packaged by Helicobacter pylori outer membrane vesicles attenuate IL-8 secretion in human cells. Int J Med Microbiol, 2020. 310(1): p. 151356.

13. Shen, Q., et al., Extracellular vesicles-mediated interaction within intestinal microenvironment in inflammatory bowel disease. J Adv Res, 2022. 37: p. 221–233.

14. Hill-Burns, E.M., et al., Parkinson’s disease and Parkinson’s disease medications have distinct signatures of the gut microbiome. Mov Disord, 2017. 32(5): p. 739–749.

15. Romano, S., et al., Meta-analysis of the Parkinson’s disease gut microbiome suggests alterations linked to intestinal inflammation. NPJ Parkinsons Dis, 2021. 7(1): p. 27.

16. Li, C., et al., Host miR-129-5p reverses effects of ginsenoside Rg1 on morphine reward possibly mediated by changes in B. vulgatus and serotonin metabolism in hippocampus. Gut Microbes, 2023. 15(2): p. 2254946.

17. Liu, S., et al., The Host Shapes the Gut Microbiota via Fecal MicroRNA. Cell Host Microbe, 2016. 19(1): p. 32–43.

18. Liu, S., et al., Oral Administration of miR-30d from Feces of MS Patients Suppresses MS-like Symptoms in Mice by Expanding Akkermansia muciniphila. Cell Host Microbe, 2019. 26(6): p. 779–794 e8.

19. Santos, A.A., et al., Host miRNA-21 promotes liver dysfunction by targeting small intestinal Lactobacillus in mice. Gut Microbes, 2020. 12(1): p. 1–18.

20. Zhao, L., et al., Colon specific delivery of miR-155 inhibitor alleviates estrogen deficiency related phenotype via microbiota remodeling. Drug Deliv, 2022. 29(1): p. 2610–2620.

21. Proano, A.C., et al., Gut Microbiota and Its Repercussion in Parkinson’s Disease: A Systematic Review in Occidental Patients. Neurol Int, 2023. 15(2): p. 750–763.

22. Schwechheimer, C. and M.J. Kuehn, Outer-membrane vesicles from Gram-negative bacteria: biogenesis and functions. Nat Rev Microbiol, 2015. 13(10): p. 605–19.

23. Dauros-Singorenko, P., et al., The functional RNA cargo of bacterial membrane vesicles. FEMS Microbiol Lett, 2018. 365(5).

24. Gerstner, N., et al., GeneTrail 3: advanced high-throughput enrichment analysis. Nucleic Acids Res, 2020. 48(W1): p. W515–W520.

25. Gardiner, C., et al., Techniques used for the isolation and characterization of extracellular vesicles: results of a worldwide survey. J Extracell Vesicles, 2016. 5: p. 32945.

26. Lehrich, B.M., Y. Liang, and M.S. Fiandaca, Foetal bovine serum influence on in vitro extracellular vesicle analyses. J Extracell Vesicles, 2021. 10(3): p. e12061.

27. Thery, C., et al., Isolation and characterization of exosomes from cell culture supernatants and biological fluids. Curr Protoc Cell Biol, 2006. Chapter 3: p. Unit 3 22.

28. Lehrich, B.M., et al., Fetal Bovine Serum-Derived Extracellular Vesicles Persist within Vesicle-Depleted Culture Media. Int J Mol Sci, 2018. 19(11).

29. Vallabhaneni, K.C., et al., Extracellular vesicles from bone marrow mesenchymal stem/stromal cells transport tumor regulatory microRNA, proteins, and metabolites. Oncotarget, 2015. 6(7): p. 4953–67.

30. Ramos, Y., S. Sansone, and D.K. Morales, Sugarcoating it: Enterococcal polysaccharides as key modulators of host-pathogen interactions. PLoS Pathog, 2021. 17(9): p. e1009822.

31. Lu, Y.C., W.C. Yeh, and P.S. Ohashi, LPS/TLR4 signal transduction pathway. Cytokine, 2008. 42(2): p. 145–151.

32. Dauros-Singorenko, P., et al., Effect of the Extracellular Vesicle RNA Cargo From Uropathogenic Escherichia coli on Bladder Cells. Front Mol Biosci, 2020. 7: p. 580913.

33. Rishik, S., et al., miRNATissueAtlas 2025: an update to the uniformly processed and annotated human and mouse non-coding RNA tissue atlas. Nucleic Acids Res, 2025. 53(D1): p. D129–D137.

34. Bradley, D.E., A function of Pseudomonas aeruginosa PAO polar pili: twitching motility. Can J Microbiol, 1980. 26(2): p. 146–54.

35. Kaiser, D., Social gliding is correlated with the presence of pili in Myxococcus xanthus. Proc Natl Acad Sci U S A, 1979. 76(11): p. 5952–6.

36. Piepenbrink, K.H., DNA Uptake by Type IV Filaments. Front Mol Biosci, 2019. 6: p. 1.

37. Chapot-Chartier, M.P. and S. Kulakauskas, Cell wall structure and function in lactic acid bacteria. Microb Cell Fact, 2014. 13 Suppl 1(Suppl 1): p. S9.

38. Krause, A.L., T.P. Stinear, and I.R. Monk, Barriers to genetic manipulation of Enterococci: Current Approaches and Future Directions. FEMS Microbiol Rev, 2022. 46(6).

39. Tan, C.H., et al., Lipid-Polymer Hybrid Nanoparticles Enhance the Potency of Ampicillin against Enterococcus faecalis in a Protozoa Infection Model. ACS Infect Dis, 2021. 7(6): p. 1607–1618.

40. Ou, F., et al., Rapid and cost-effective evaluation of bacterial viability using fluorescence spectroscopy. Anal Bioanal Chem, 2019. 411(16): p. 3653–3663.

41. Bhardwaj, V., et al., snakePipes: facilitating flexible, scalable and integrative epigenomic analysis. Bioinformatics, 2019. 35(22): p. 4757–4759.

42. Ewels, P., et al., MultiQC: summarize analysis results for multiple tools and samples in a single report. Bioinformatics, 2016. 32(19): p. 3047–8.

43. Dobin, A., et al., STAR: ultrafast universal RNA-seq aligner. Bioinformatics, 2013. 29(1): p. 15–21.

44. Tarasov, A., et al., Sambamba: fast processing of NGS alignment formats. Bioinformatics, 2015. 31(12): p. 2032–4.

45. Liao, Y., G.K. Smyth, and W. Shi, featureCounts: an efficient general purpose program for assigning sequence reads to genomic features. Bioinformatics, 2014. 30(7): p. 923–30.

46. Love, M.I., W. Huber, and S. Anders, Moderated estimation of fold change and dispersion for RNA-seq data with DESeq2. Genome Biol, 2014. 15(12): p. 550.

47. Fehlmann, T., et al., miRMaster 2.0: multi-species non-coding RNA sequencing analyses at scale. Nucleic Acids Res, 2021. 49(W1): p. W397–W408.

48. Griffiths-Jones, S., et al., miRBase: microRNA sequences, targets and gene nomenclature. Nucleic Acids Res, 2006. 34(Database issue): p. D140–4.

49. Molder, F., et al., Sustainable data analysis with Snakemake. F1000Res, 2021. 10: p. 33.

50. Hunter, J.D., Matplotlib: A 2D Graphics Environment. Computing in Science & Engineering, 2007. 9(3): p. 90–95.

51. Waskom, M., *seaborn: statistical data visualization*. Journal of Open Source Software, 2021. 6(60).

52. Wickham, H., ggplot2: Elegant Graphics for Data Analysis. Springer-Verlag New York, 2016.

