## Supplementary_Figures for "Extracellular vesicles and their RNA cargo facilitate bidirectional cross-kingdom communication between human and bacterial cells"

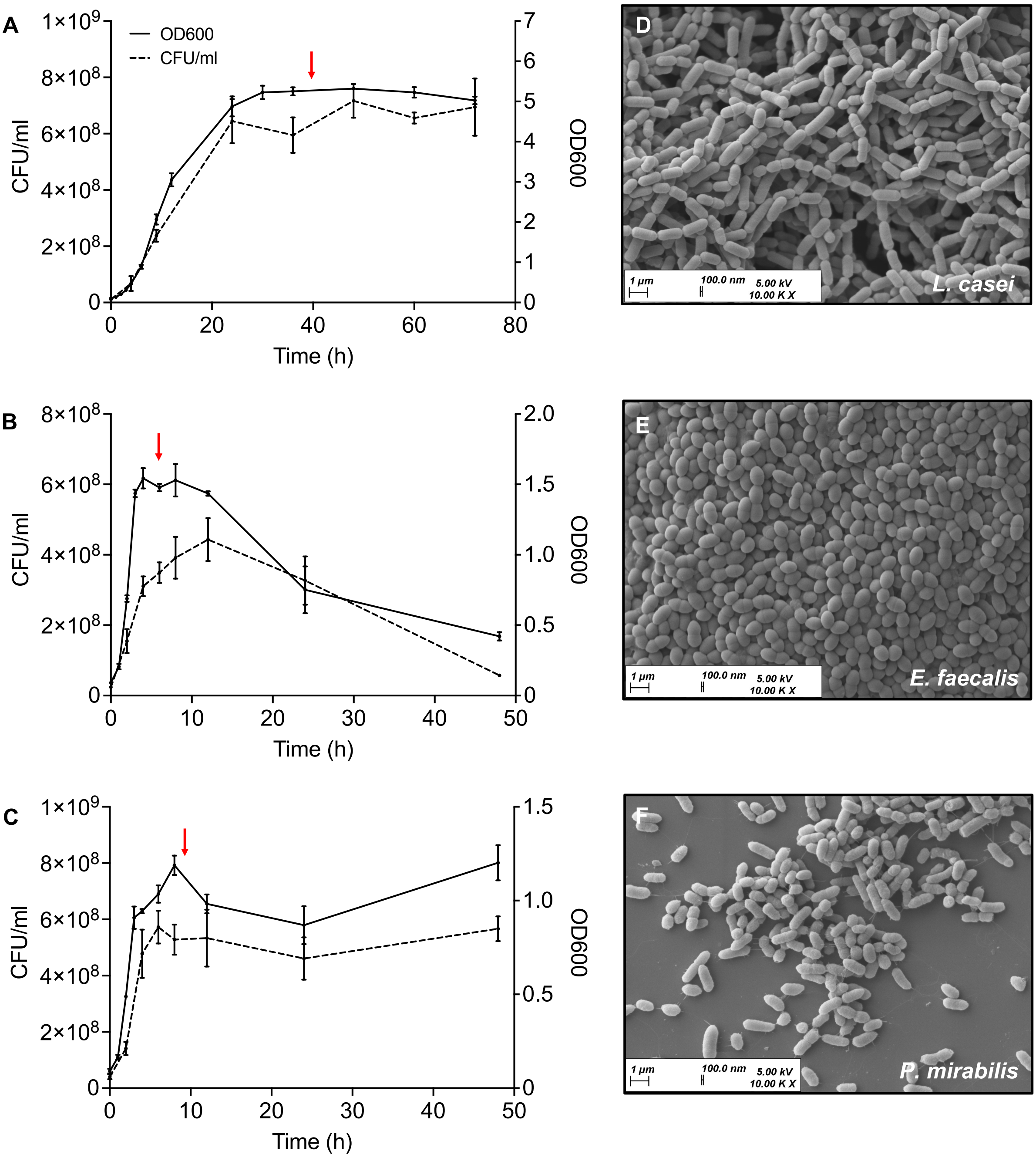


**Figure S1: Analysis of bacterial growth and morphology.**
(**A-C**) Growth of *L. casei* (**A**), *E. faecalis* (**B**), and *P. mirabilis* (**C**) over a period of 72 h or 48 h at 37 °C. Results are presented as mean ± standard deviation (SD) from 3-4 independent experiments. The red arrows mark the timepoints where the supernatant for BEV isolation was collected (*L. casei:* 40 h, *E. faecalis:* 6 h, *P. mirabilis:* 9 h) (**D-F**) Scanning electron microscopic images of *L. casei* (**D**), *E. faecalis* (**E**), and *P. mirabilis* (**F**). Bacteria were grown for 40 h (*L. casei*) or 12 h (*E. faecalis* or *P. mirabilis*) at 37 °C. Afterwards, the bacteria were fixed, washed and sputter coated. Images were taken at 5kV and 10,000x magnification.

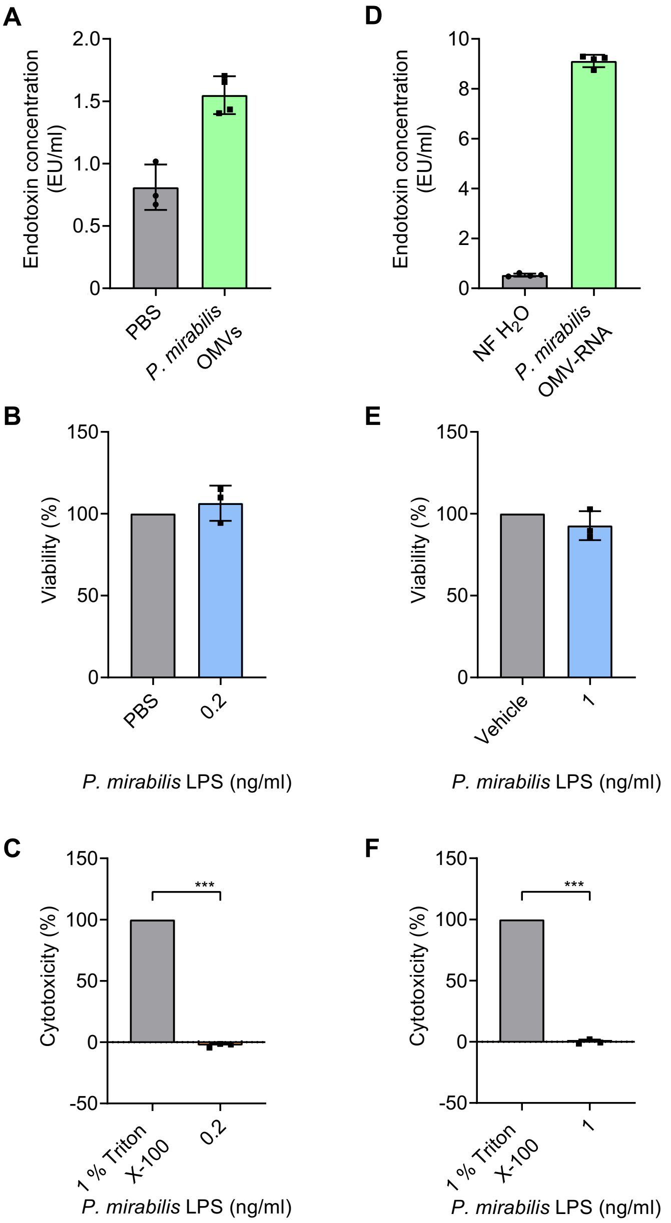


**Figure 2: Quantification of LPS concentration and effects of LPS on the viability of Caco-2 cells.** (**A-C**) Quantification and effects of LPS in *P. mirabilis* OMVs. (**A**) Endotoxin concentration in (EU/ml) of purified *P. mirabilis* OMVs in comparison to PBS. (**B and C**) Measurement of cell viability and cytotoxicity after incubation of Caco-2 cells with *P. mirabilis* LPS for 24 h. 1 % Triton X-100 was used as dead control, whereas PBS was used as live control. (**D-F**) Quantification and effects of LPS in *P. mirabilis* OMV-RNA. (**D**) Endotoxin concentration in (EU/ml) of purified *P. mirabilis* OMV-RNA in comparison to NF $H_2O$ (Nuclease-free water). (**E and F**) Measurement of cell viability and cytotoxicity after transfection of Caco-2 cells with *P. mirabilis* LPS for 24 h. 1 % Triton X-100 was used as dead control, whereas Lipofectamine™ 3000 transfection reagent mixed with nuclease-free water (vehicle) was used as live control. All results are presented as mean ± standard deviation (SD) from 3-4 independent experiments. For all viability and cytotoxicity assays, a two-tailed unpaired t-test was conducted to determine statistical significance ($p < 0.001$ ***).

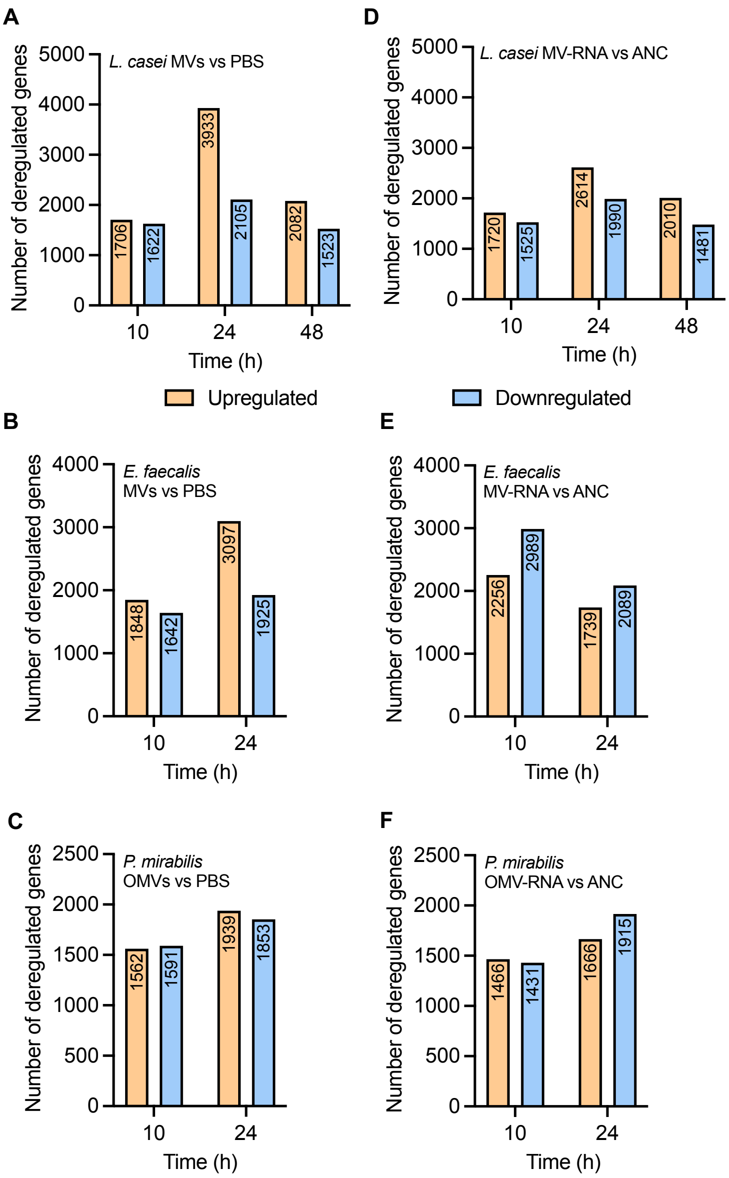


**Figure S3: Number of deregulated genes after incubation of Caco-2 cells with BEVs or transfection with BEV-RNA.**

(**A-F**) Number of deregulated genes after incubation of Caco-2 cells with $1.5 \times 10^{11}$ *L. casei* MVs (**A**), $7.5 \times 10^{9}$ *E. faecalis* MVs (**B**), or $6.0 \times 10^{9}$ *P. mirabilis* OMVs (**C**), or after transfection of Caco-2 cells with 100 ng *L. casei* MV-RNA (**D**), 5 ng *E. faecalis* MV-RNA (**E**), or 2 ng *P. mirabilis* OMV-RNA (**F**).

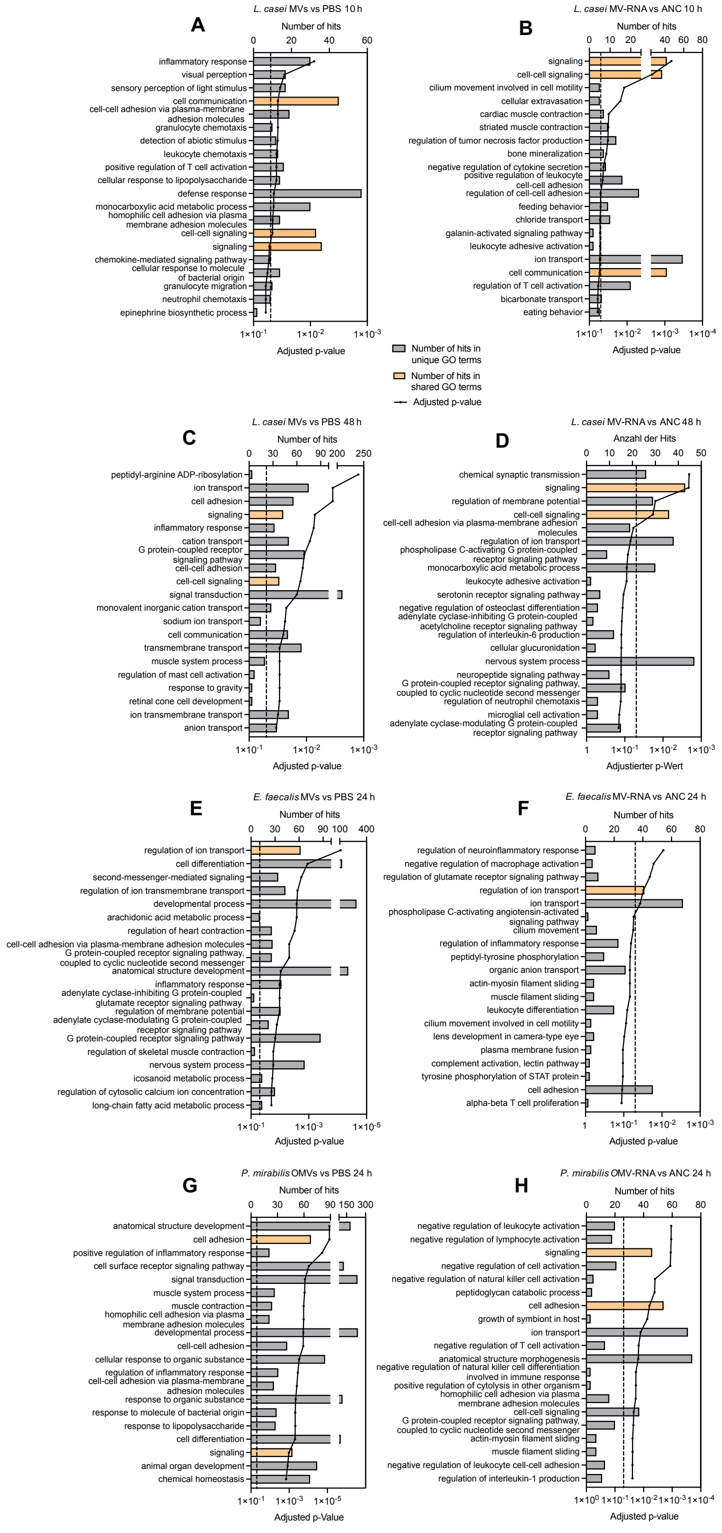


**Figure S4: Enrichment of genes in specific biological processes after incubation of Caco-2 cells with BEVs or transfection with BEV-RNA.**

(**A-H**) Caco-2 cells have been incubated with BEVs (*L. casei* MVs: $1.5x10^{11}$, *E. faecalis* MVs: $7.5x10^{9}$, *P. mirabilis* OMVs: $6.0x10^{9}$) or transfected with BEV-RNA (*L. casei* MV-RNA: 100 ng, *E. faecalis* MV-RNA: 5 ng, *P. mirabilis* OMV-RNA: 2 ng) for 10 h, 24 h, or 48 h. Differentially expressed genes were used to perform an over-representation analysis (ORA) using GeneTrail3.2. Shown are the top 20 Gene Ontology (GO) - biological process in which the differentially expressed genes were enriched in. The dashed line marks the significance level of p = 0.05. Biological processes that showed an enrichment after incubation with BEVs and transfection of BEV-RNA derived from the same bacteria at the same timepoint are highlighted in orange.

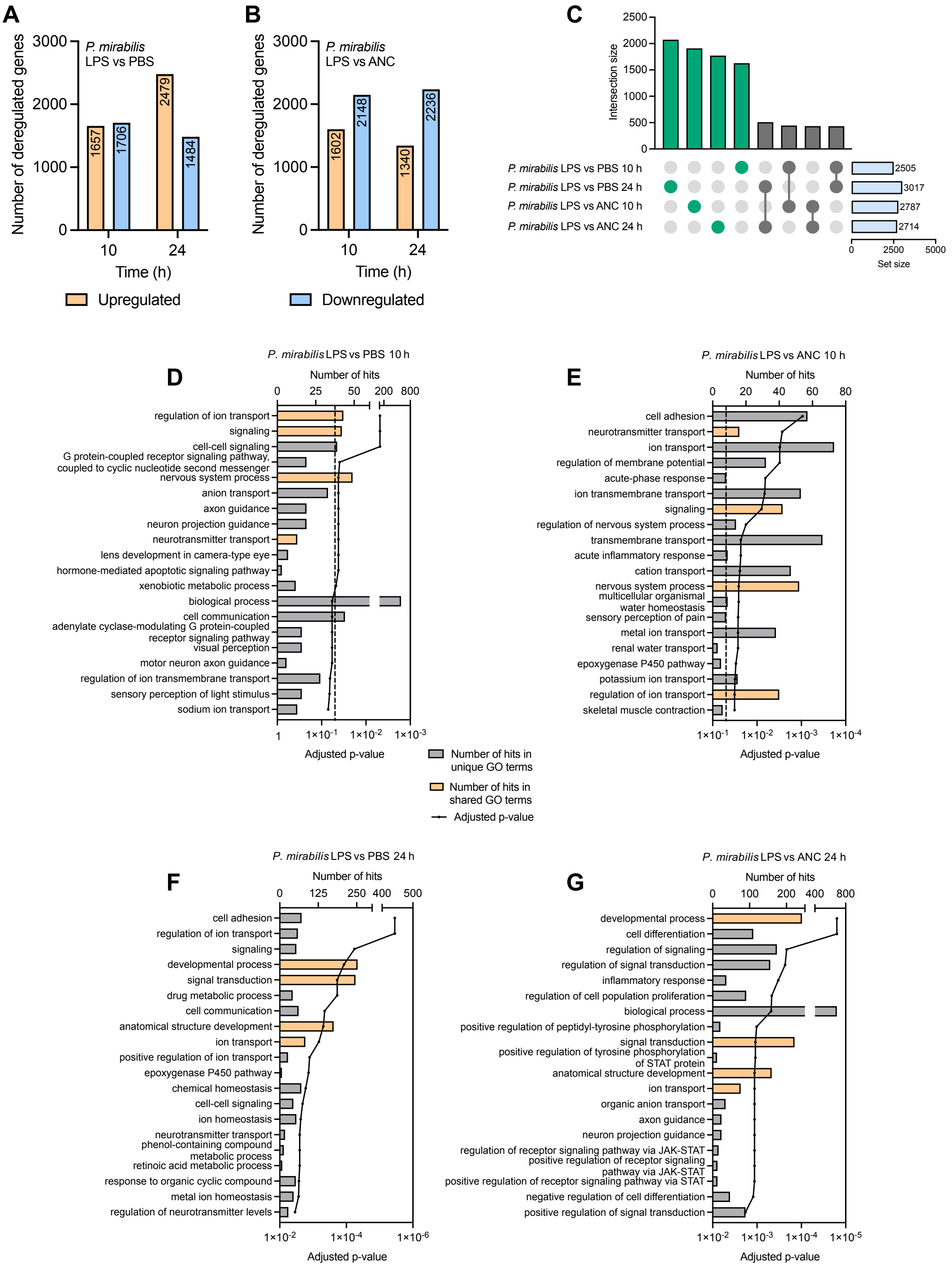


**Figure S5: Changes in the gene expression of Caco-2 cells after incubation or transfection with *P. mirabilis* LPS.**

(**A and B**) Number of deregulated genes after incubation of Caco-2 cells with 0.2 ng/µl *P. mirabilis* LPS or transfection with 1 ng/µl *P. mirabilis* LPS. (**C**) Upset plot highlights the number of differentially expressed genes between the different time points and treatments. (**D-G**) Differentially expressed genes were used to perform an over-representation analysis (ORA) using GeneTrail3.2. Shown are the top 20 Gene Ontology (GO) - biological process in which the differentially expressed genes were enriched in. The dashed line marks the significance level of p = 0.05. Biological processes that showed an enrichment after incubation with BEVs and transfection of BEV-RNA derived from the same bacteria at the same timepoint are highlighted in orange.

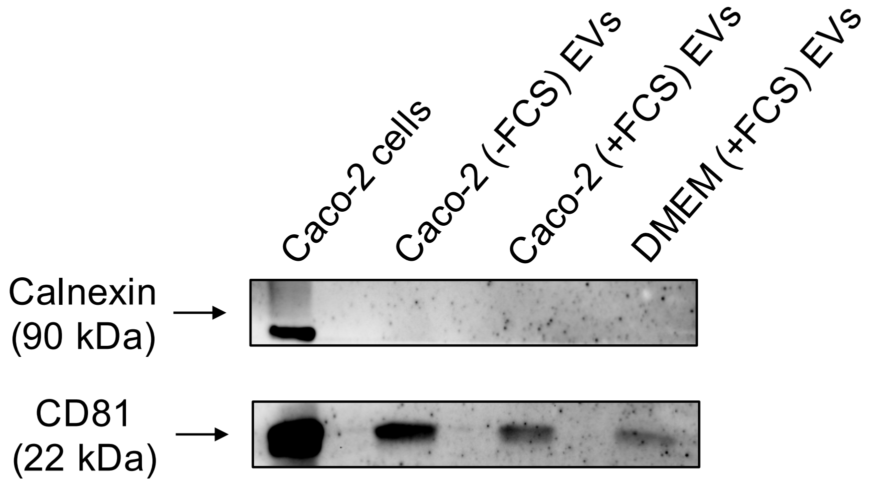


**Figure S6: Detection of cellular and EV marker using Western Blot analysis.**
Representative Western blot for calnexin (cellular marker) and CD81 (EV marker) in purified EVs from different conditions and Caco-2 cells.

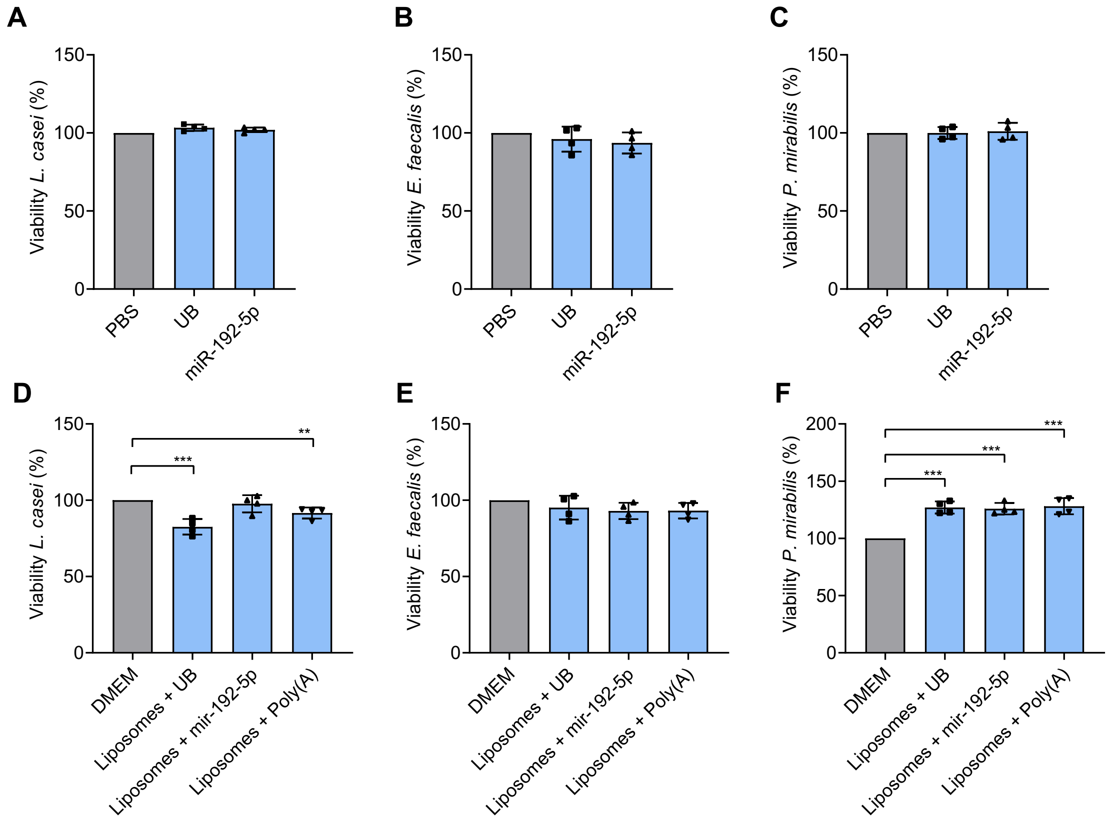


**Figure S7: Effects of miR-192-5p on the viability of bacteria.**
(**A-C**) Viability of bacteria in the presence of free miR-192-5p. Percentage of live *L. casei* (**A**), *E. faecalis* (**B**) and *P. mirabilis* (**C**) after cultivation in the presence of 4 µM free synthetic miR-192-5p or 6 µl 1 x siMAX Universal Buffer (UB) for 40 h (*L. casei*) or 24 h (*E. faecalis* and P. *mirabilis*) at 37 °C. PBS was used as control. (**D-F**) Viability of bacteria in the presence of liposome-packaged miR-192-5p. Percentage of live *L. casei* (**D**), *E. faecalis* (**E**) and *P. mirabilis* (**F**) after cultivation in the presence of 4 µM liposome-packaged synthetic miR-192-5p, liposomes in combination with 6 µl 1 x siMAX Universal Buffer ( UB) or 8 µg liposome-packaged Poly(A) for 40 h (*L. casei*) or 24 h (*E. faecalis* and P. *mirabilis*) at 37 °C. DMEM was used as control. All results are presented as mean ± standard deviation (SD) from 4 independent experiments. For all experiments, a two-tailed unpaired t-test was conducted to determine statistical significance ($p \leq 0.05$ *, $p \leq 0.01$ **, $p \leq 0.001$ ***).
